# Differential coupling of adult-born granule cells to parvalbumin and somatostatin interneurons

**DOI:** 10.1101/598615

**Authors:** Ayelén I. Groisman, Sung M. Yang, Alejandro F. Schinder

## Abstract

The dentate gyrus of the hippocampus is dominated by a strong GABAergic tone that maintains sparse levels of activity. Adult neurogenesis disrupts this balance through the continuous addition of new granule cells (GCs) that display high excitability while develop and connect within the preexisting host circuit. The dynamics of the connectivity map for developing GCs in the local inhibitory networks remains unknown. We used optogenetics to study afferent and efferent synaptogenesis between new GCs and GABAergic interneurons expressing parvalbumin (PV-INs) and somatostatin (SST-INs). Inputs from PV-INs targeted the soma and remained immature until they grew abruptly in >4-week-old GCs. This transition was accelerated by exposure to enriched environment. Inputs from SST-INs were dendritic and developed slowly until reaching maturity by 8 weeks. Synaptic outputs from GCs onto PV-INs matured faster than those onto SST-INs, but also required several weeks. In the mature dentate network, PV-INs exerted an efficient control of GC spiking and were involved in both feedforward and feedback loops, a mechanism that would favor lateral inhibition and sparse coding. Our results reveal a long-lasting transition where adult-born neurons remain poorly coupled to inhibition, which might enable a parallel streaming channel from the entorhinal cortex to CA3 pyramidal cells.

## INTRODUCTION

Activity-dependent changes in synaptic connectivity are thought to underlie learning and long-term memory storage. In the dentate gyrus of the mammalian hippocampus, including humans, plasticity also involves the generation of new neurons that develop, integrate and contribute to information processing (Goncalves et al., 2016; Mongiat and Schinder, 2011; Moreno-Jimenez et al., 2019; van Praag et al., 2002; Zhang et al., 2016). Adult-born granule cells (GCs) play differential roles in processing spatial information and resolve specific behavioral demands, such as the identification of subtle contextual cues required for spatial discrimination (Clelland et al., 2009; Kropff et al., 2015; Nakashiba et al., 2012; Sahay et al., 2011). They are also relevant for behavioral responses to fear and stress (Anacker and Hen, 2017; Anacker et al., 2018; Guo et al., 2018). Moreover, impaired adult neurogenesis has been associated to cognitive dysfunctions that are commonly found in patients with psychiatric disorders (Kang et al., 2016). Developing granule cells (GCs) interact dynamically with the preexisting network, changing their intrinsic and synaptic characteristics as they grow towards morphological and functional maturation (Mongiat and Schinder, 2011). With time, GABA signaling switches from excitation to inhibition, excitability decreases and excitatory inputs grow in number, reaching mature characteristics after 6 to 8 weeks (Ge et al., 2007a; Laplagne et al., 2006; Temprana et al., 2015). GCs undergo a transient period of high excitability and plasticity due to their reduced inhibition, which is consequence of the weak strength and slow kinetics of GABAergic postsynaptic responses (Marin-Burgin et al., 2012). Understanding the rules that guide integration of new GCs in the host networks is essential for harnessing adult neurogenesis as a mechanism of brain plasticity in health and disease.

GABAergic interneurons (INs) control the excitation/inhibition balance of principal cells in all regions of the mammalian brain, which is critical to achieve an overall network homeostasis (Isaacson and Scanziani, 2011). GABAergic circuits encompass distinct neuronal subtypes, whose functional relevance in different brain areas remains to be determined. Ivy/neurogliaform INs contact GCs from early developmental stages and coordinate the network activity with different IN populations (Markwardt et al., 2011). Parvalbumin-(PV) and somatostatin-expressing (SST) cells represent two major classes of INs in the hippocampus (Hosp et al., 2014; Kepecs and Fishell, 2014). PV-INs represent ~30% of the population and their axons target perisomatic compartments of postsynaptic neurons (Freund, 2003; Freund and Buzsaki, 1996). They contribute to the synchronization of principal cell activity and the generation of network oscillations (Bartos et al., 2007). In the dentate gyrus, they display the highest degree of connectivity compared to other INs (Espinoza et al., 2018). SST-INs represent ~50% of GABAergic INs and primarily target dendritic compartments in postsynaptic cells. They are a heterogenous group that provides local and long-range inhibition and are implicated in hippocampal-prefrontal synchrony during spatial working memory (Abbas et al., 2018; Yuan et al., 2017). GABAergic INs contact adult-born GCs before the onset of glutamatergic synaptogenesis, and these initial connections play critical roles in shaping development and integration of new GCs (Alvarez et al., 2016; Espósito et al., 2005; Ge et al., 2006; Overstreet Wadiche et al., 2005; Song et al., 2013). Yet, the developmental time-course of GABAergic synaptogenesis and the precise contribution of PV-INs and SST-INs to inhibition in new GCs remain unclear.

In this study, we show that PV-INs and SST-INs establish functional synapses onto new GCs at early development but these connections require several weeks to reach functional maturation, enabling a mechanism for long-lasting remodeling of local circuits. Contacts from PV-INs develop faster, and synaptic transmission during the period of high excitability is modulated by experience. Outputs from GCs onto PV-INs also mature earlier than those onto SST-INs. Interestingly, while both IN populations establish feedback loops in the GCL, feedforward loops from the perforant path onto the GCL are primarily mediated by PV-INs. Our results reveal that adult neurogenesis produces a neuronal population that remains apart from the inhibitory tone dominating dentate gyrus activity, enabling a parallel channel for input processing that is also involved in long-lasting circuit reorganization.

## RESULTS

### GABAergic synaptogenesis onto developing GCs

To investigate how inhibition becomes established in new GCs, we characterized the connectivity between developing GCs and two of the main types of dentate gyrus interneurons; PV-INs and SST-INs. PV^Cre^ and SST^Cre^ mice were utilized to express channelrhodopsin-2 (ChR2) in either interneuron population by crossing them with CAG^floxStopChR2EYFP^ mice (Ai32) (Hippenmeyer et al., 2005; Madisen et al., 2012; Taniguchi et al., 2011). Retroviral labeling was used to express red fluorescent protein (RV-RFP) in newly generated GCs of the same mice. PV^Cre^; CAG^floxStopChR2EYFP^ mice labeled a homogeneous neuronal population that expressed the calcium buffer parvalbumin. Their bodies were localized primarily in the GCL, their axons spread along the GCL, and they displayed high spiking frequency (>100 Hz), typical of GABAergic basket interneurons (**Fig. 1A,B**; **Fig. S1**). SST^Cre^; CAG^floxStopChR2EYFP^ mice labeled neurons that expressed the neuropeptide somatostatin, localized primarily in the hilar region, and displaying variable spiking patterns, corresponding to a heterogeneous population of GABAergic interneurons (**Fig. 1E,F**; **Fig. S2**).

**Figure 1.**
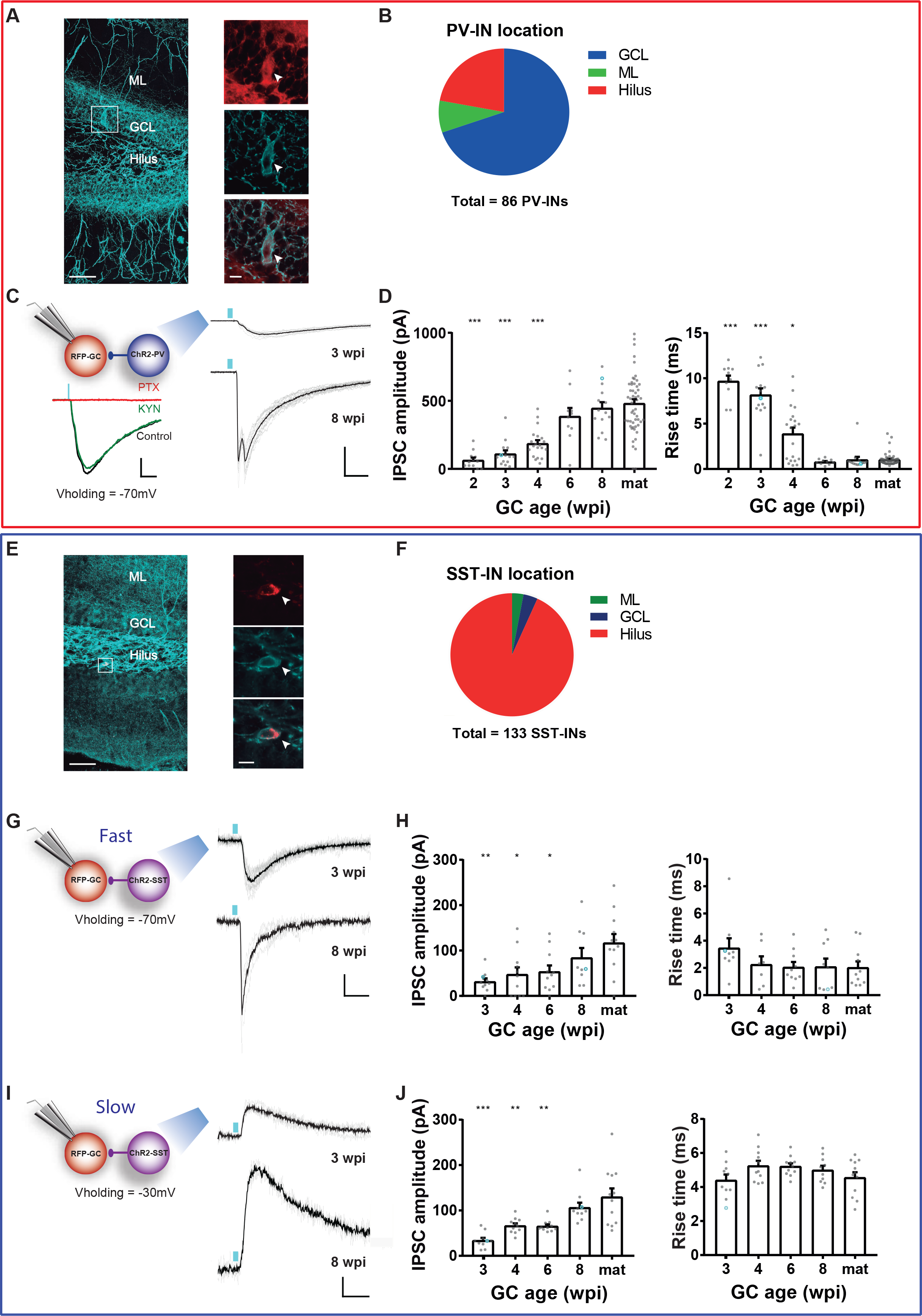
GABAergic synaptogenesis onto developing GCs. (A) Confocal image of a 60 μm-thick hippocampal section depicting PV-INs in a PV^Cre^;CAG^floxStopChR2-EGFP^ mouse. (GCL: granule cell layer; ML: molecular layer). Scale bar, 50 μm. Insets show single optical planes of PV-INs (soma indicated by the arrow) displaying immunolabeling for PV (red), expression of ChR2-EYFP (blue), and their overlay (bottom). Scale bar, 20 μm. (B) Distribution of cell body localization in different areas of the DG. (C) *Left panel,* experimental scheme of laser-mediated stimulation of PV-INs combined with IPSCs recordings in adult-born GCs, and example traces from a 4-wpi GC showing blockade by PTX (100 μM, red) but not by KYN (6 mM, green). Scale bars, 50 pA, 20 ms. *Right panel,* IPSCs elicited by laser pulses (0.2 ms, blue marks) delivered at low frequency (0.07 Hz), recorded from GCs at 3 and 8 wpi. Traces depict individual sweeps (gray) and their average (black). Scale bars, 200 pA, 20 ms. (D) IPSC peak amplitude and rise time for different GC ages. Dots correspond to individual neurons. Blue circles correspond to example traces shown in (C). (E) Confocal image depicting SST-INs in a SST^Cre^;CAG^floxStopChR2-EGFP^ mouse. Scale bar, 50 μm. Insets show single optical planes of SST-INs (soma indicated by the arrow) displaying immunolabeling for SST (red), ChR2-EYFP (blue), and their overlay. Scale bar, 10 μm. (F) Cell body localization in different areas of the DG. (G-J) Laser stimulation of SST-INs evoked IPSCs with different kinetics and reversal potentials. Recordings performed at V_h_=−70 mV (G,H) elicited fast IPSCs, whereas traces obtained at V_h_=−30 mV (I,J) were slower. Scale bars 20 pA, 10 ms. Sample sizes, >11 neurons from >4 mice (PV-INs), and >9 neurons in >2 mice (SST-INs). Statistical comparisons were done using one-way ANOVA followed by *post hoc* Bonferroni’s test for multiple comparisons against mature condition, with *p*<0.05 (*), *p*<0.01 (**) and *p*<0.001 (***).

Stereotaxic surgery was performed in 6–7-week-old mice to deliver a RV-RFP in a cohort of new GCs. ChR2-expressing INs were reliably activated using brief laser pulses (0.2 ms), which elicited spikes with short onset latency (**Fig. S3A-C, Fig. S4A-C**). Whole cell recordings were performed on RFP-GCs in acute slices at 2-8 weeks post injection (wpi). Laser stimulation of PV-INs elicited inhibitory postsynaptic currents (IPSCs) in RFP-GCs that were completely abolished by the GABA_A_ receptor antagonist picrotoxin (100 μM), but were not affected by the ionotropic glutamate receptor blocker kynurenic acid (KYN, 6 mM) (**Fig. 1C**; **Fig. S3D,E**). Together with the fast IPSC onset, these data reveal that PV-INs make monosynaptic GABAergic contacts onto adult-born CGs (**Fig. S3H**). Activation of ChR2-PVs reliably elicited IPSCs already in 2 wpi GCs, but responses displayed small amplitude and slow kinetics, typical of immature synapses (**Fig. S3F,G**). As GC development progressed, the amplitude of postsynaptic responses increased and kinetics became substantially faster, as revealed by the reduction of half-width and rise time, particularly in the window between 4 and 6 wpi (**Fig. 1D**; **Fig. S3E-J**). In fact, 4 weeks can be visualized as a transition point with two split populations where some GCs display slow rise time and others have already became fast. Remarkably, while synapse formation from PV-INs to GCs was initiated early in development (before 2 wpi), synaptic maturation was only apparent at >6 wpi, when IPSCs reached fastest kinetics and maximal amplitude. Interestingly, the age-dependent growth in IPSC amplitude was mainly due to an increased quantal size rather than changes in the number of synaptic contacts; no differences were found in the number of functional synapses between young and mature GCs, measured as the ratio between IPSC in saturation and unitary IPSC amplitude (**Fig. S3K-M**). These results demonstrate a slow age-dependent maturation of the PV-IN to GC synapse.

ChR2-SSTs also formed functional monosynaptic contacts onto new GCs as early as 2-3 wpi (**Fig. 1G-J**; **Fig. S4F-I,L,M**). At these developmental stages, activation of SST-INs reliably elicited IPSCs with small amplitude and slow kinetics (**Fig. S4H-O**). Two types of responses were distinguished based on kinetics, coefficient of variation of the amplitude (**Fig. S4D,E**) and reversal potential; one slow component observed at a depolarized membrane potential, and one fast that was visualized at hyperpolarized potentials. To determine their nature, their amplitude and kinetics were measured by holding the membrane at the reversal potential of the alternate component. Both responses displayed age-dependent increase in IPSC amplitude (**Fig. 1G-J**). However, the kinetic features for both components remained fundamentally unchanged through GC maturation (**Fig. 1H,J**; **Fig. S4H-O**). Finally, mature synaptic properties were only observed in GCs at >8 wpi. Together, these results show that new GCs receive monosynaptic GABAergic inputs from PV-INs and SST-INs early in development, and both connections become gradually strengthened along maturation, acquiring mature synaptic properties at 6 to 8 weeks of age.

### Differential subcellular localization of synapses formed by PV-INs and SST-INs

In whole-cell recordings, the intracellular Cl^−^ concentration ([Cl^−^]_i_) near the soma is imposed by the recording patch pipette, whereas Cl^−^ transporters in distal dendritic compartments can overcome the pipette load and maintain physiological levels of [Cl^−^]_i_ (Khirug et al., 2005). This gradient in [Cl^−^]_i_ results in differences in the reversal potential of GABA-mediated currents along the somato-dendritic axis (Laplagne et al., 2007; Pearce, 1993). To reveal the subcellular localization of the PV-IN to GC synapse, we monitored the reversal potential of optogenetically activated currents by means of whole-cell recordings under conditions that resulted in an equilibrium potential for [Cl^−^]_i_ of −30 mV at perisomatic compartments. Extracellular stimulation of GABAergic axons in the outer molecular layer (OML) was used to activate distal dendritic inputs (Laplagne et al., 2007)(**Fig. 2A-E**). Thus, activation of PV-INs elicited fast IPSCs with depolarized reversal potential for all neuronal ages, suggesting that synaptic localization was close to the recording compartment (soma) and remained stable throughout GC development. In contrast, OML stimulation evoked slow IPScs with hyperpolarized reversal potentials as expected for the physiological [Cl^−^]_i_ (~−70 mV), suggesting that they originated in the dendritic compartment, located distally from the recording site. In fact, fast-inward and slow-outward IPSCs were simultaneously observed at intermediate membrane potentials (V_h_=−50 mV) when ChR2-PVs and OML stimulation were combined (**Fig. 2B**). These results demonstrate a perisomatic origin for PV-IN-mediated IPSCs at all GC ages.

**Figure 2.**
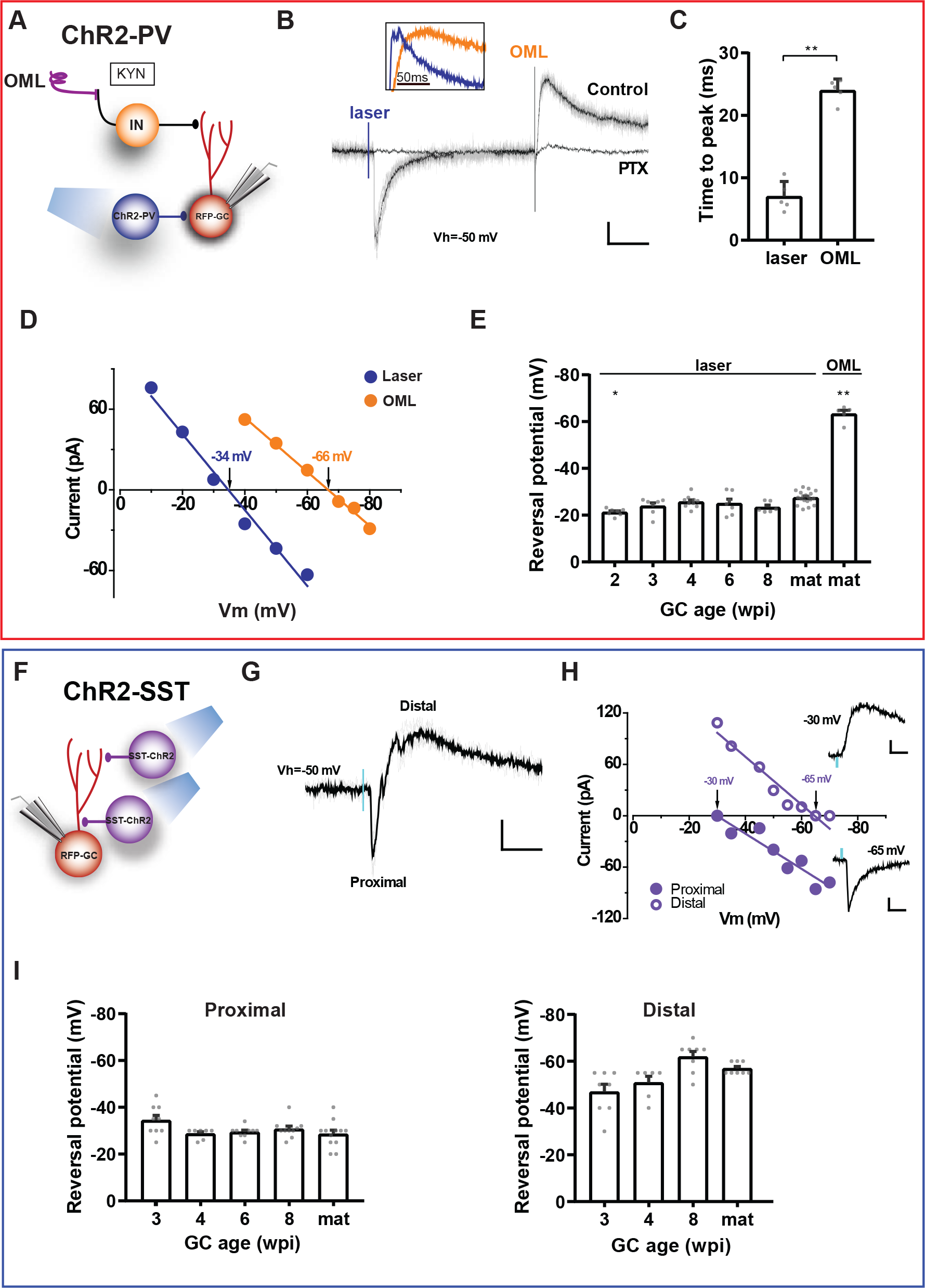
Differential localization of synapses formed by PV-INs and SST-INs. (A) Experimental scheme to compare responses of adult-born GC elicited by laser stimulation of ChR2-PVs versus electrical stimulation in the OML. (B) IPSCs elicited by laser pulses (0.2 ms, blue mark) and by electrical stimulation in the OML. All responses were blocked by PTX (100 μM). Scale bars, 100 ms, 10 pA. The inset shows normalized IPSCs to highlight the difference in kinetic. (C) Time to peak of evoked responses in mature GCs by laser and OML stimulation. Statistical comparison was done using Mann-Whitney’s test, with n=5 neurons (5 slices). (D) I-V curves for responses shown in (B), with reversal potentials indicated by the arrows. (E) Reversal potential for different GC stages. Statistical comparisons were done using Kruskal-Wallis’ test followed by *post hoc* Dunn’s multiple comparisons against mature GCs with laser stimulation. N=6 to 18 cells. (F) Experimental scheme of laser-mediated stimulation of ChR2-SSTs to assess their subcellular target location. (G) IPSCs elicited by single laser pulses. Recordings performed at −50 mV show bi-phasic currents corresponding to proximal (early onset) and distal (delayed) components. Scale bars, 20 ms, 20 pA. (H) I-V curves for both responses shown in (G). Reversal potentials are indicated by arrows. Insets show isolated IPSCs recorded at the reversal potential of the other component. Scale bars, 10 ms, 20 pA. (I) Reversal potentials for different GC ages. N = 6-8 cells (IPSC-distal) and 5-12 cells (IPSC-proximal). Statistical comparisons were done using Kruskal-Wallis test followed by *post hoc* Dunn’s multiple comparisons against mature GCs. *p*<0.05 (*) and *p*<0.01 (**).

Stimulation of ChR2-SSTs elicited mixed inward and outward IPSCs in adult-born GCs held at −50 mV (**Fig. 2F-H**), arising from synaptic responses originated in compartments with different distances to the soma. Indeed, the fast current exhibited a depolarized reversal potential (~−30 mV), consistent with a proximal localization, whereas the slow current reversed at more negative potentials (up to ~−60 mV), suggesting a distal contact. Proximal IPSCs maintained similar values for reversal potential through GC development, while distal IPSCs showed a subtle but progressive hyperpolarization, consistent with the observation that control of [Cl^−^]_i_ homeostasis improves during neuronal development (Khirug et al., 2005)(**Fig. 2I**). We conclude that ChR2-SSTs establish functional synapses onto new GCs with distinct proximal and distal localizations.

### Short-term plasticity of GABAergic responses

During normal behavior, networks of principal neurons and interneurons exhibit complex patterns of activation and undergo spiking discharges in a wide range of frequencies. Under these conditions, synapses are subject to short- and long-lasting activity-dependent modifications of synaptic transmission (Hsu et al., 2016; Lee et al., 2016; Pardi et al., 2015). To investigate how repetitive activity impinges on postsynaptic responses in developing GABAergic synapses, ChR2-PVs or ChR2-SSTs were stimulated by brief trains (5 laser pulses at 20 Hz) and whole-cell recordings were performed in developing GCs. Responses to ChR2-PVs stimulation displayed short-term depression that became more pronounced as GCs matured (**Fig. 3A-D)**. These results reveal changes in presynaptic release machinery along synaptic maturation.

**Figure 3.**
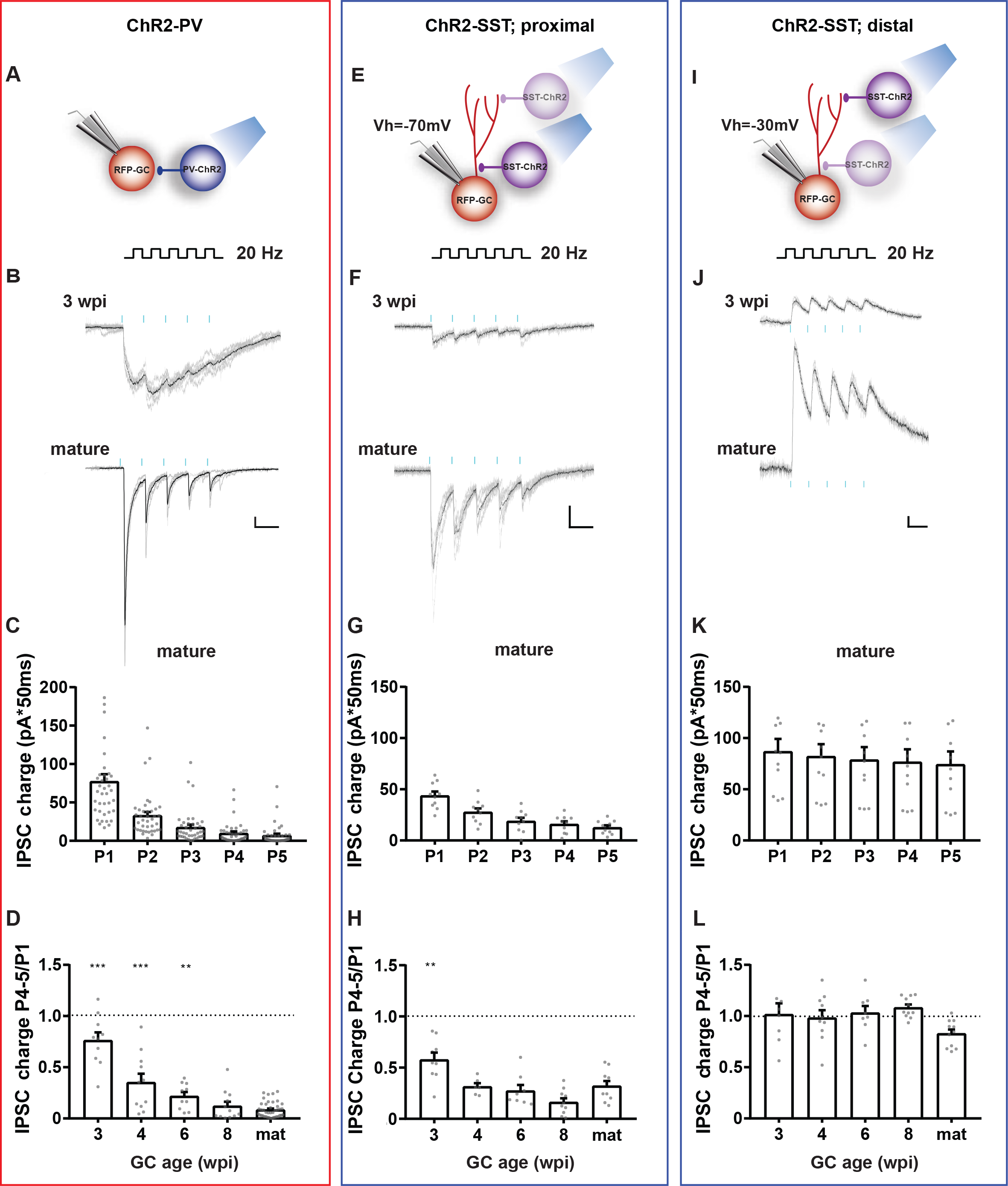
Short-term plasticity of IPSCs. (A) Experimental scheme for recording postsynaptic responses elicited by repetitive stimulation of PV-INs (5 pulses, 0.2 ms, 20 Hz). (B) IPSCs recorded from GCs at different ages in response to trains delivered at 0.07 Hz (blue marks). Traces depict all sweeps (gray) and their average (black). Scale bars, 50 ms, 20 pA. (C) IPSC charge for individual pulses of the train (P1-P5), recorded in mature GCs. (D) IPSC charge for pulses 4 and 5 (P4-5) normalized to the charge in the first pulse, for the indicated ages of postsynaptic GCs. (E-H) Proximal postsynaptic responses elicited by repetitive stimulation of SST-INs (GC V_holding_ = −70 mV). (I-L) Distal responses elicited by repetitive stimulation of SST-INs (GC V_holding_ = −30 mV), recorded in the same set of neurons shown in (*E-H*). Statistical comparisons were done using one-way ANOVA followed by *post hoc* Bonferroni’s test for multiple comparisons against the mature group. Sample sizes (presented as GCs/mice): 8-41/4-20 for PV-INs; 6-11/2-5 for SST-INs, with *p*<0.01 (**), and *p*<0.001 (***).

SST-IN to GC synapses of proximal and distal locations were discriminated by their reversal potential and their responses upon repetitive stimulation were analyzed separately. Activation of ChR2-SSTs by brief trains (20 Hz) induced a marked short-term depression in proximal IPSCs, which became more pronounced in more mature GCs (**Fig. 3E-H**). In contrast, distal IPSCs showed stable pulse amplitudes along the train and no signs of depression for any of the GC ages (**Fig. 3I-L**). These results further support the conclusion that proximal and distal responses evoked by SST-INs belong to functionally different synapses.

### GABAergic interneurons control activity in the granule cell layer

The impact of PV-INs and SST-INs on spiking activity of the granule cell layer (GCL) was monitored in field recordings of excitatory postsynaptic potentials (fEPSPs) evoked by stimulation of the medial performant path (mPP) (**Fig. 4A**). In these recordings, the area of the population spike (pop-spike) is proportional to the number of active GCs, and the fEPSP slope reflects the strength of the synaptic input. Paired activation of ChR2-PVs with mPP stimulation modulated the fEPSP response; increasing laser power recruited more PV-INs, which resulted in a progressive and reliable reduction of the pop-spike (**Fig. 4B**). SST-INs were also able to control GCL recruitment, but they exerted a smaller effect over the pop-spike than PV-INs. Maximum inhibitory effects were found when PV-INs or SST-INs and mPP axons were simultaneously stimulated (**Fig. 4C-F**). In addition, inhibition of the pop-spike by PV-IN activation was more efficient and acted over a broader time interval compared to SST-INs, in concordance with their larger IPSCs and perisomatic targeting. These data demonstrate that both types of INs can modulate spiking in the GCL, although control by PV-INs is more reliable, probably due to the somatic localization of their synapses.

**Figure 4.**
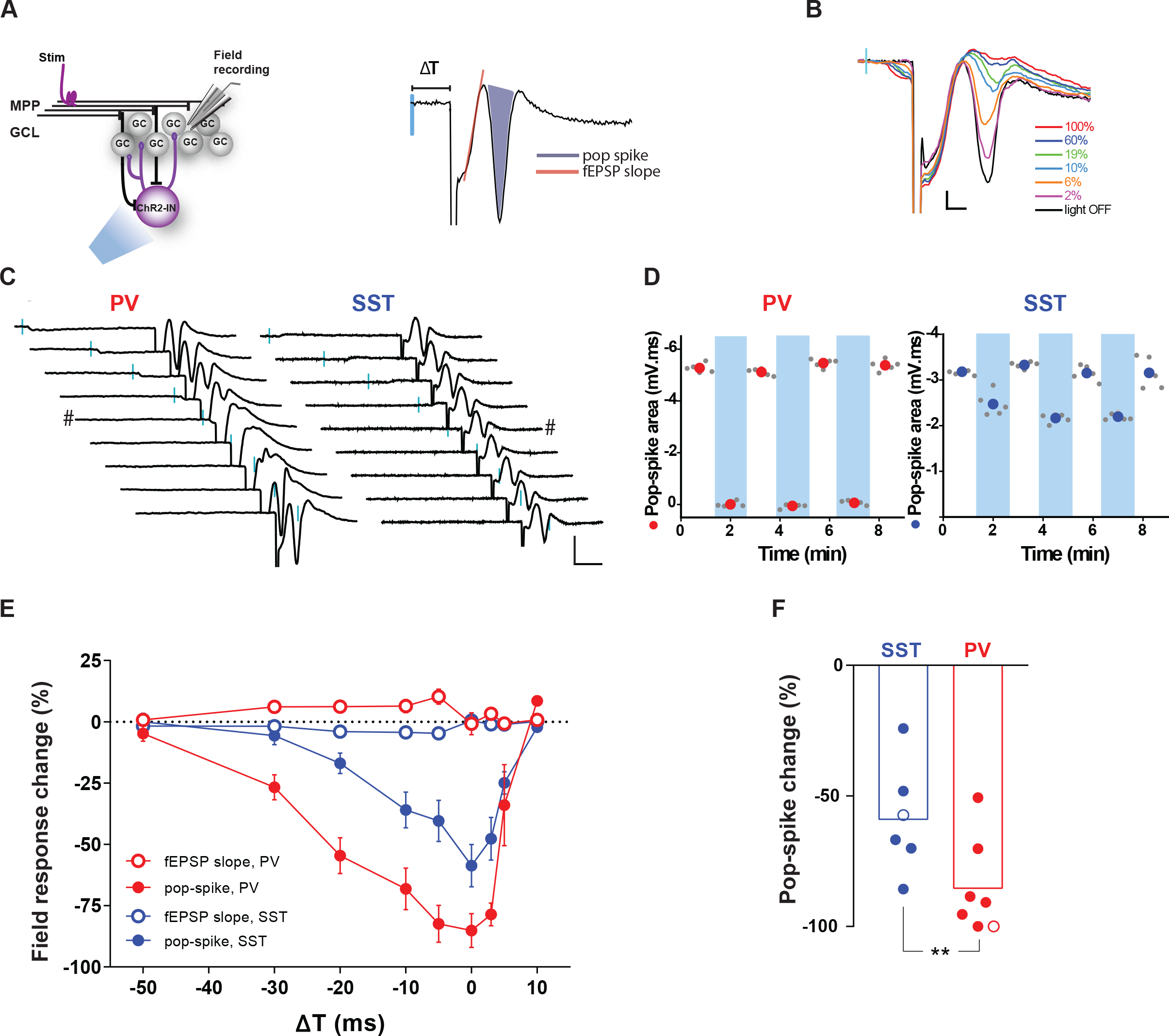
PV-INs and SST-INs control GCL spiking. (A) Experimental design for field potential recordings. The example trace on the right illustrates fEPSP changes elicited by activation of mPP fibers combined with laser activation of ChR2-expressing INs with variable delays (ΔT). The shaded area is proportional to the number of spiking neurons in the GCL (pop-spike). (B) Activation of ChR2-PVs with increasing laser power intensity (ΔT=−5 ms) abolishes pop-spikes triggered by mPP stimulation. Scale bars: 0.2 mV, 2 ms. (C) Subsequent fEPSP recordings for progressive delays (from −50 to +10 ms) between mPP stimulation and laser activation of ChR2-PVs (left) or ChR2-SSTs (right). Scale bars, 10 ms, 1 mV. (D) Pop-spike areas produced by low frequency stimulation (0.07 Hz) of mPP alone (white columns) or paired with preceding laser pulses (ΔT=−5 ms; blue bars). Colored circles represent mean values. (E) Laser-induced change of field responses defined as 100*(fEPSP_mPP_ −fEPSP_mPP+laser_)/fEPSP_mPP_. Data obtained from 7 slices/6 animals (PV-INs) and 6 slices/3 animals (SST-INs). (F) Pop-spike change by optogenetic activation of the indicated INs paired simultaneously to electrical stimulation (ΔT=0). Hollow circles correspond to example traces indicated by # in (*C*). Statistical comparisons were done using Mann-Whitney’s test, with *p*<0.01 (**).

### Functional synaptogenesis of GC outputs onto local interneurons

To map the networks of GABAergic interneurons activated by adult-born GCs, we used retroviruses to selectively express ChR2-GFP in cohorts of new GCs (ChR2-GCs) at different stages of development (3 to 11 wpi). Reliable activation of ChR2-GCs was achieved by laser stimulation (1-ms pulses; **Fig. S5A-C**), allowing the study of synaptic responses in INs. PV^Cre^;CAG^floxStop-tdTomato^ and SST^Cre^;CAG^floxStop-tdTomato^ were used to label PV-INs and SST-INs, respectively, and perform whole-cell recordings of excitatory postsynaptic currents (EPSCs; **Fig. 5A-J**). Activation of developing GCs elicited glutamatergic excitatory postsynaptic currents in both PV-INs and SST-INs, but no functional connections were detected before 4–6 wpi (**Fig. S5D-Q**). When responses occurred, they displayed short onset latency (**Fig. S5H,O)** and were blocked by KYN (not shown), indicating that these glutamatergic connections are monosynaptic. At early ages, GCs elicited a large proportion of transmission failures. As neurons became more mature, the proportion of failures decreased to reach a plateau that occurred at 6 weeks for PV-INs and >8 weeks for SST-INs (**Fig. 5E,J**).

**Figure 5.**
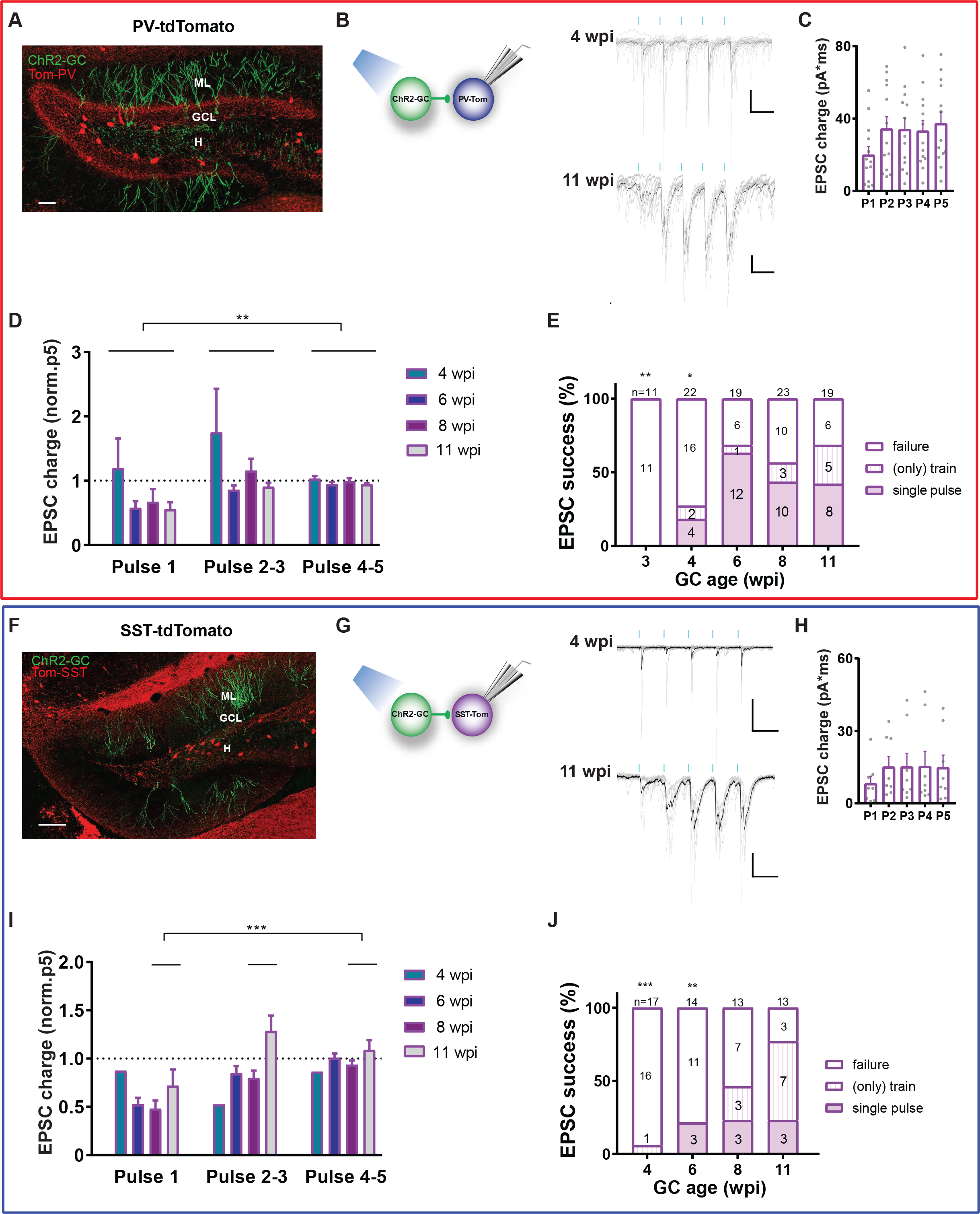
Short-term plasticity of EPSCs evoked by new GCs onto PV-INs and SST-INs. (A) Confocal image of a 60 μm-thick hippocampal section depicting PV-INs (red) and 6-week-old GCs expressing GFP-ChR2 (green) in a PV^Cre^;CAG^floxStoptdTom^ mouse (ML: molecular layer; H: hilus). Scale bar, 100 μm. (B) EPSCs obtained from PV-INs evoked by laser stimulation of ChR2-GCs at the indicated ages (5-pulse trains at 0.07 Hz, 1 ms, 20 Hz; blue marks). Traces depict individual sweeps (gray) and their average (black). Scale bars, 50 ms, 50 pA. (C) EPSC charge for individual pulses (P1-P5) delivered to 11-wpi ChR2-GCs. Dots correspond to individual neurons. (D) EPSC charge normalized to the fifth pulse (P5). (E) Proportion of INs that displayed EPSC upon activation of new GCs by single pulses or trains, at the indicated ages. Given the short-term facilitation of this synapse, in some cases the EPSC was elicited by train stimulation but not by single pulses. Total numbers of recorded PV-INs are shown on top of each column (n = 5 to 9 mice). Responsive INs to different protocols are displayed inside the columns. (F) Confocal image depicting SST-INs (red) and 6-week-old GCs expressing GFP-ChR2 (green) in a SST^Cre^;CAG^floxStoptdTom^ mouse. Scale bar, 100 μm. (G) EPSCs obtained from SST-INs evoked by laser stimulation of ChR2-GCs at the indicated ages. Traces depict individual sweeps (gray) and their average (black). Recordings were done as described in (*B*). Scale bars, 50 ms, 50 pA. (H) EPSC charge for individual pulses delivered to 11-wpi ChR2-GCs. (I) EPSC charge normalized to P5. (J) Proportion of INs that displayed EPSC upon activation of new GCs at the indicated ages (n = 2 to 7 mice). Statistical comparisons were done using 2-way ANOVA test followed by *post hoc* Tukey’s (D, I), and Fisher’s exact test agains the 11 wpi group (E,J), with *p*<0.05 (*), *p*<0.01 (**) and *p*<0.001 (***).

GCs activation in awake behaving rodents can cover a broad range of discharge activity. To better characterize the physiological significance of GC to IN connections, we delivered brief trains of laser stimulation (5 pulses at 20 Hz) onto ChR2-GCs. In contrast to the depression that was typically observed in IPSCs (**Fig. 3**), EPSCs displayed strong facilitation at all developmental stages in both PV-INs and SST-INs (**Fig. 5B-D,G-I**). Facilitation resulted in decreased failures in synaptic transmission along subsequent pulses within a train, suggesting that repetitive firing in GCs is more likely to activate GABAergic INs than individual spikes. In fact, train stimulation revealed connections that remained silent when assessed by individual stimuli (**Fig. 5E,J**). Taking into account the EPSC success rate, which represents the likelihood of finding functional synaptic connections, our data indicate that immature GCs are reliable in establishing connections onto PV-INs, while SST-INs receive sparse inputs.

### Contribution of PV-INs and SST-INs to inhibitory loops

Dentate gyrus INs participate in feedforward (FFI) and feedback (FBI) inhibitory microcircuits, with functional impact in both the GCL and CA3. To dissect the participation of PV-INs and SST-INs in those inhibitory loops, we designed an experiment that allowed both an efficient recruitment of IN spiking and the assessment of feedback and feedforward pathways.

We thus combined whole-cell recordings in PV- or SST-INs with simultaneous field recordings in the GCL, and measured responses to electrical stimulation of the mPP to a level that evoked a reliable pop-spike (~50 % of maximum response). When recording from PV-INs, mPP activation typically elicited two action potentials, one occurring before the peak of the pop-spike and another occurring after a brief delay (**Fig. 6A-C**). This sequence suggests that the first spike was evoked directly by mPP activation, while the second one was evoked by activation of the heterogeneous GC population (including both mature and developing neurons). To test this possibility, we used DCG-IV, an agonist of group II metabotropic glutamate receptors that reduces release probability in mossy fiber terminals and in mPP terminals in the GCL (Kamiya et al., 1996; Macek et al., 1996). DCG-IV reduced the amplitude of the fEPSP response, eliminating the pop-spike, which in turn abolished the second PV-IN spike without altering the first one (**Fig. 6D,E**). Subsequent application of KYN blocked the first spike. Together, these results demonstrate that the same individual PV-INs are recruited by mPP axons that activate a feedforward inhibitory loop and by GCs that recruit a feedback loop in the GCL.

**Figure 6.**
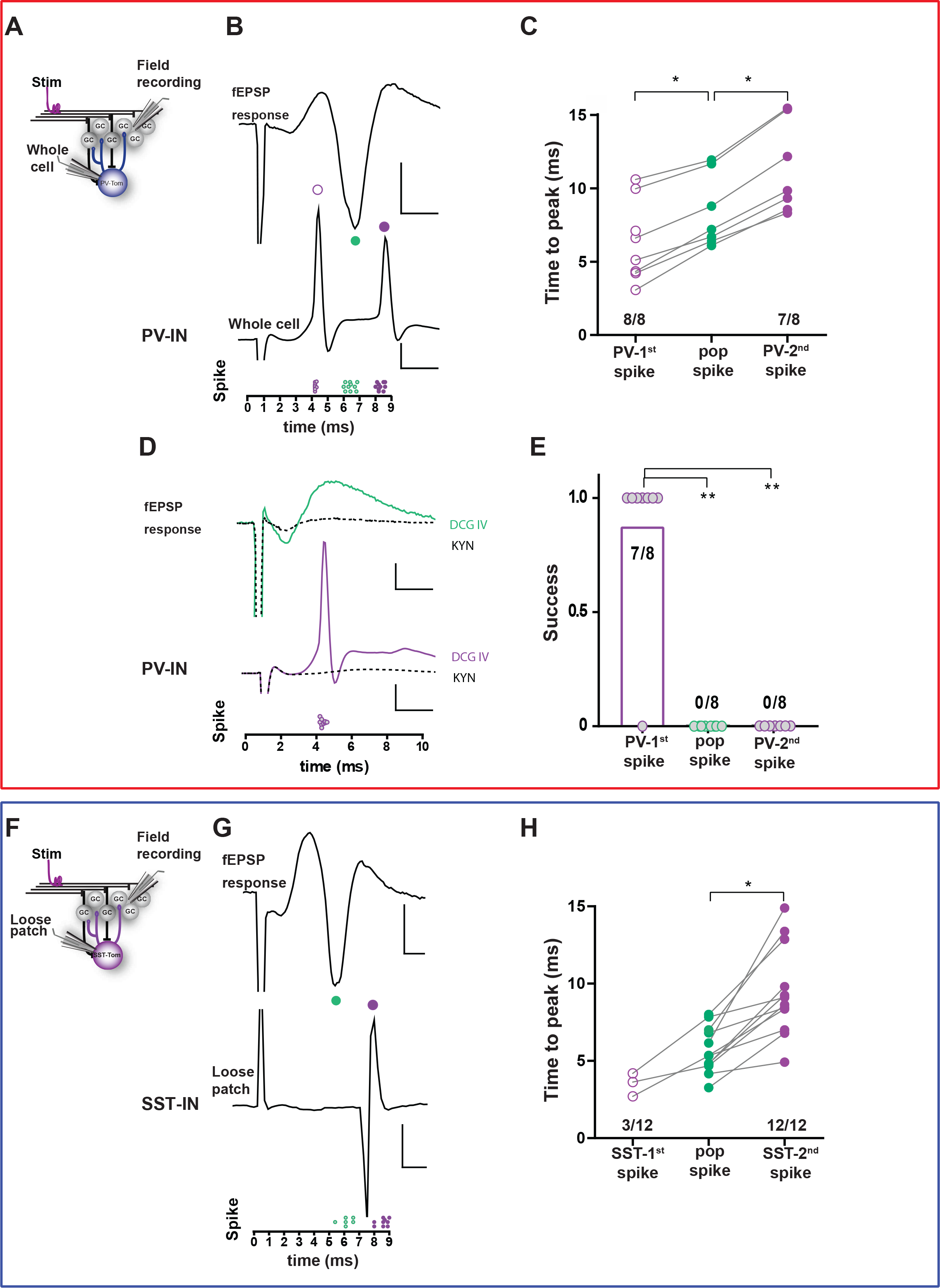
Differential recruitment of PV-INs and SST-INs by local excitatory networks. (A) Experimental scheme: simultaneous recordings of fEPSP in the GCL and membrane potential in PV-INs were carried out in response to mPP stimulation. (B) Example fEPSP (top) and whole-cell recordings in a PV-IN (middle), together with measurements of time to peak for spikes. Scale bars, 2 ms, 1 mV (top), 20 mV (bottom). (C) Delay to spike for all individual experiments. N=8 PV-INs, 7 slices, 5 mice. Statistical comparisons were done using Friedman test followed by Wilcoxon matched-pairs signed rank test, with *p*<0.05 (*). (D) DCG-IV (green trace) prevents GCs pop spike and, consequently, the second PV-IN spike triggered by GC activity (purple). KYN (6 mM) suppressed all spikes (black dotted lines). Scale bars, 2 ms, 0.5 mV (top), 20 mV (bottom). (E) Rate of success to evoke spikes in presence of DCG-IV. Number of cases (positive/total) are shown. Statistical comparisons were done using Fisher’s exact test, with *p*<0.01 (**). (F) Experimental scheme. (G) Example fEPSP (top) and loose patch recording in a SST-IN (bottom), together with measurements of time to peak for spikes. Scale bars, 2 ms, 1 mV (top), 100 pA (middle). (H) Delay to spike for all individual experiments N=12 SST-INs, 10 slices, 10 mice. Statistical comparison was done using Wilcoxon matched-pairs signed rank test, with *p*<0.01 (**).

In contrast, the same assay showed that SST-INs were primarily recruited to trigger action potentials after the pop-spike (12/12 neurons), with only a small proportion activated before (3/12, **Fig. 6F-H**). Thus, SST-INs mainly participate in feedback inhibition, while their participation in the feedforward inhibitory loop is scarce. Together, these results demonstrate that cortical activity reaching the dentate gyrus through mPP axons recruit feedforward inhibition through PV-INs that exert tight control over GC spiking (Fig. 4). In turn, GCs activate a feedback inhibitory loop by PV-INs, now acting in concert with SST-INs to provide finely tuned activation of the GCL.

### Modulation of perisomatic inhibition by experience

We have previously shown that experience in enriched environment (EE) can promote early development of newly generated GCs, with PV-INs acting as key transducers from behavior to local circuit rearrangement (Alvarez et al., 2016). We now investigated whether experience can also influence synaptic connections of PV-INs onto more developed GCs that may already be involved in information processing. RV-GFP was delivered into PV^Cre^; CAG^floxStopChR2EYFP^ mice that were then exposed to regular cages or switched to EE for 2 weeks. Synaptic transmission in response was analyzed at 4 wpi (**Fig. 7A,B**). Single laser pulses elicited IPSCs of similar amplitude in both conditions, but responses obtained from EE mice displayed faster kinetics, consistent with a more mature synapse (**Fig. 7C-F**). This difference was more evident when ChR2-PVs were stimulated with 50-Hz trains. In control mice, GCs presented individual responses to repetitive pulses that accumulate along the train, finalizing with a slow decay after the last stimulus. In contrast, signals from mice exposed to EE displayed faster kinetics, resulting in a progressive depression of the synaptic response (**Fig. 7G-I**). These results demonstrate that transmission in this developing synapse is sensitive to experience in a manner that favors a mature behavior.

**Figure 7.**
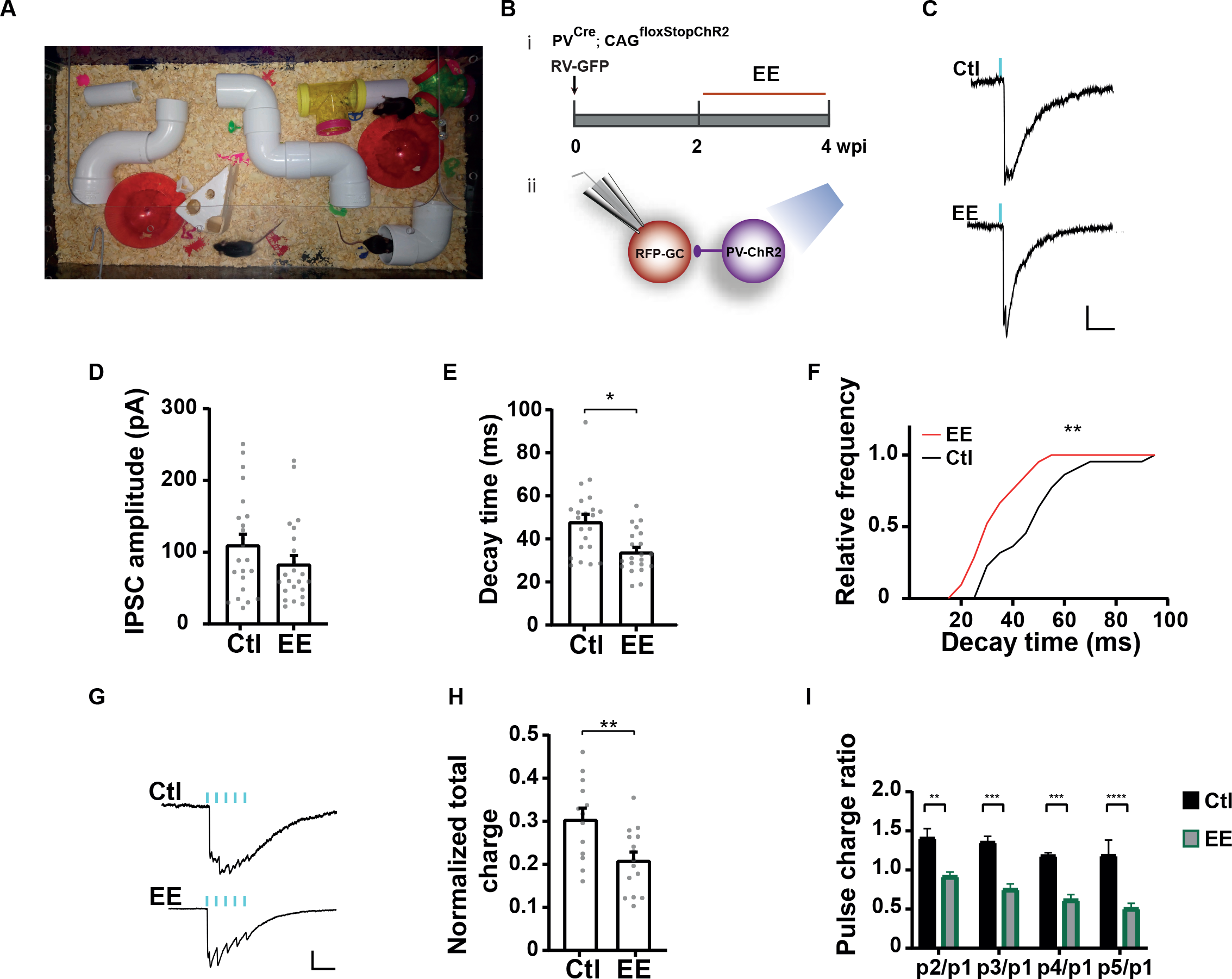
Modulation of perisomatic GABAergic inhibition by EE. (A) Image of the EE cage. (B) Experimental design. (i) RV-GFP was delivered to young adult mice housed in control or EE exposure from 2 to 4 wpi (red segment). (ii) Laser-mediated stimulation of PV-INs evokes IPSCs in 4wpi GCs. (B) Representative traces of averaged IPSCs elicited by laser pulses (0.2 ms, blue marks) delivered at 0.07 Hz, recorded from GCs at 4 wpi obtained from control or EE mice. Scale bars, 50 pA, 50 ms. (D-E) Amplitude and kinetics of individual evoked IPSCs. Gray dots represent individual cells. (G-I) IPSCs evoked in response to 50 Hz trains of laser pulses (delivered at 0.035 Hz). (G) Example average traces. Scale bars, 50 ms, 50 pA. (H) Charge of the entire IPSC over 340 ms, normalized to the peak amplitude. (I) Charge of individual pulses within the train normalized to the 1^st^ peak. Sample sizes were 13-23 cells from >5 animals for both control and EE conditions. Statistical comparisons were done using Mann-Whitney test with (*) *p*<0.05 (D), *t*-test with (**) *p* = 0.0013 (E) and (**) *p*=0.0055 (H), Kolmogorov-Smirnov test (**) p = 0.0097 (F) and two-way Anova followed by Tukey’s multiple comparisons (I): (**) *p*<0.01, (***) p<0.001, (****) p<0.0001.

## DISCUSSION

The function that neurons acquire in a given circuit depend on their intrinsic properties, relevant for signal integration, and their connections, which determine network dynamics. It has been proposed that developing GCs play unique functional roles in DG computation (Clelland et al., 2009; Kim et al., 2012; Kropff et al., 2015). We found a slow development of connectivity between new GCs and GABAergic INs, which conveys immature GCs the property to behave as computational modules with rules that vary along development and are different to those of mature GCs. PV-IN synapses onto new GCs are formed early (Song et al., 2013) but, as shown here, synaptic responses mature during several weeks, becoming increasingly stronger and faster. The slow IPSC kinetics exhibited by young synapses may well explain the hyperexcited neuronal behavior described previously in 4-week old GCs (Marin-Burgin et al., 2012; Pardi et al., 2015). When IN activity occurs in bursts instead of single spikes (train stimulation in our experiments), synaptic maturation results in a reduction of the integration time, transforming slow responses with sustained postsynaptic currents (in 3 to 4-week-old GCs), into faster postsynaptic responses with pronounced depression (8-week-old GCs). We found that this synapse is sensitive to behavioral stimuli; experience in EE accelerated the transition to the fast responsiveness typical of a mature synapse. Because PV-INs are involved both in FBI and FFI, a change in kinetics like the one reported here would produce substantial changes at the network level (Hu et al., 2014).

Optical stimulation of SST-INs generated two types of responses that differed in kinetics and reversal potential when measured in mature GCs. We observed a fast response with a depolarized reversal potential that revealed proximal localization, and a slower response with a more hyperpolarized reversal potential that corresponded to dendritic (distal) distribution. ChR2-SST stimulation using 20 Hz trains resulted in substantial synaptic depression in proximal responses but stable amplitude in distal synaptic currents, which strengthened the idea of separate proximal and distal responses. It is unclear whether they correspond to different axons of the same INs or different INs altogether. Electrophysiological characterization of intrinsic properties revealed four groups of SST-INs with distinctive spiking patterns and input resistance (Fig. S2). In this context, we speculate that proximal and distal synapses derive from individual populations of SST-INs that target different cellular compartments. In fact, two subtypes of SST-INs were recently reported in the DG and might underlie the responses observed here; hilar-perforant-path-associated INs with axon fibers in the molecular layer that make distal synapses, and INs with axons in the hilus that provide perisomatic inhibition (Yuan et al., 2017).

Activation of mature GCs recruit feedback loops that limit activation of the GCL through lateral inhibition (Espinoza et al., 2018; Temprana et al., 2015). They also recruit mossy cells that activate a range of interneuron cell types, with a preference for basket cells (Scharfman, 2018). As shown here using direct optogenetic activation, both PV-INs and SST-INs can limit activation of the GCL and could mediate the FBI triggered by GCs, although PV-INs are more efficient (**Fig. 4**), probably due to the localization and strength of their output contacts and their high degree of network connectivity (Espinoza et al., 2018). We performed two experiments to monitor FBI in the network. First, optogenetic activation of adult-born GCs revealed that both PV-INs and SST-INs are direct targets of new GCs with increasing synaptic strength as they approach maturation. Second, activation of PV-INs and SST-INs occurred following the GCL pop-spike elicited by stimulation of mPP axons, and did not occur when the pop-spike was blocked by DGC IV (**Fig. 6**). Together, these results show that the FBI loop that controls activity of the GCL involves both INs. As expected, the feedforward loop was activated by mPP stimulation independently of the presence of the pop-spike and primarily involved PV-INs rather than SST-INs, consistent with a higher efficacy of the connectivity of mPP axons towards basket PV-INs (Hsu et al., 2016).

Jonas and collaborators have recently obtained a thorough map of the dentate gyrus network assessing the connectivity between mature GCs and different types of GABAergic INs. They found that PV-INs are the most extensively connected type of GABAergic IN and, in this network, inhibition is much more abundant than excitation (Espinoza et al., 2018). They also showed that PV-INs preferably contact GCs from which they receive no input, thus favoring lateral over recurrent inhibition by about 10 fold. It was proposed that such architecture favors a winner-takes-all model in which GCs that are strongly recruited during a particular behavior will dominate activity in the dentate gyrus. This model would be compatible with pattern separation, a network computation where similar inputs are converted into non-overlapping patterns of network spiking and might be crucial for hippocampal functions that include spatial navigation and contextual discrimination (Drew et al., 2013; McAvoy et al., 2015). Interestingly, lateral inhibition would require the coincident activation of a number of excitatory inputs from GCs to reach spiking threshold of PV-INs (Espinoza et al., 2018). Our finding that the same individual PV-INs participate in feedforward and feedback inhibition suggests that excitation from mPP axons might contribute to lower the threshold for efficient activation of PV-INs by a sparse population of active GCs.

Using a simple computational model, we have proposed that adult neurogenesis may favor the acquisition of non-overlapping input spaces through the delayed coupling to inhibition of developing GCs (Kropff et al., 2015; Temprana et al., 2015). Our new results demonstrate that, during several weeks, developing GCs remain poorly coupled to the IN networks both at the input and output levels, escaping lateral inhibition and creating a parallel channel for the information flow from entorhinal cortex to CA3 (Marin-Burgin et al., 2012; Temprana et al., 2015). During this period, activity-dependent synaptic modifications might refine input and output connections required to encode relevant information on the acquired task (Ge et al., 2007b; Gu et al., 2012). As we have shown here, experience modulates this network at the level of the PV-IN synapse during a critical period of high excitability in new GCs. With time, inhibition becomes more efficient and new GCs are more sparsely activated.

## ACKNOWLEDGMENTS

We thank Verónica Piatti, the members of the A.F.S. lab and Guillermo Lanuza lab for insightful discussions. A.F.S. is investigator in the Consejo Nacional de Investigaciones Científicas y Técnicas (CONICET). S.M.Y. and A.I.G. were supported by CONICET fellowships. This work was supported by grants from the Argentine Agency for the Promotion of Science and Technology (PICT2015-3814 and PICT2016-0675), Howard Hughes Medical Institute (#55007652), and NIH (R01NS103758) to A.F.S.

## MATERIALS AND METHODS

### Animals and surgery for retroviral delivery

Genetically modified mice Pvalb^*tm1(cre)Arbr*^ mice (Hippenmeyer et al., 2005), kindly provided by S. Arber, Sst^*tm2.1(cre)Zjh/J*^ mice (Taniguchi et al., 2011), and CAG^floxStop-tdTomato^(Ai14) (B6;129S6-*Gt(ROSA)26Sor*^*tm14(CAG-tdTomato)Hze*^/J) conditional reporter line (Madisen et al., 2010), obtained from Hongkui Zeng, were crossed to generate PV^Cre^; CAG^FloxStopTom^ mice and SST^Cre^; CAG^FloxStopTom^ mice to label PV- and SST-expressing GABAergic interneurons (Tom-PV and Tom-SST), respectively. Pvalb^*tm1(cre)Arbr*^ and Sst^*tm2.1(cre)Zjh/J*^ mice were also crossed with CAG^floxStopChR2-EYFP^(Ai32) (Gt(ROSA)26Sor^*tm32(CAGCOP4*H134R/EYFP)Hze*^/J) mice from Jackson Laboratories, to generate PV^Cre^;CAG^FloxStopChR2^ and SST^Cre^; CAG^FloxStopChR2^ mice. Mice were maintained in C57Bl/6J background.

Genetically modified mice of either sex were used at 6 – 7 weeks of age, housed at 2 – 4 mice per cage. Running wheel housing started 2-3 days before surgery and continued until the day of slice preparation, to maximize the number of retrovirally transduced neurons. For surgery, mice were anesthetized (150 μg ketamine/15 μg xylazine in 10 μl saline/g), and virus (1 – 1.2 μl at 0.15 μl/min) was infused into the dorsal area of the right dentate gyrus using sterile microcapillary calibrated pipettes and stereotaxic references (coordinates from bregma: −2 mm anteroposterior, −1.5 mm lateral, −1.9 mm ventral). Experimental protocols were approved by the Institutional Animal Care and Use Committee of the Leloir Institute according to the Principles for Biomedical Research involving animals of the Council for International Organizations for Medical Sciences and provisions stated in the Guide for the Care and Use of Laboratory Animals.

### Retroviral vectors

A replication-deficient retroviral vector based on the Moloney murine leukemia virus was used to specifically transduce adult-born granule cells as done previously (Marin-Burgin et al., 2012; Piatti et al., 2011). Retroviral particles were assembled using three separate plasmids containing the capside (CMV-vsvg), viral proteins (CMV-gag/pol) and the transgenes: CAG-RFP or channelrhodopsin-2 (ChR2; Ubi-ChR2-EGFP retroviral plasmid, kindly provided by S. Ge, SUNY Stony Brook). Plasmids were transfected onto HEK 293T cells using deacylated polyethylenimine. Virus-containing supernatant was harvested 48 h after transfection and concentrated by two rounds of ultracentrifugation. Virus titer was typically ~10^5^ particles/μl. CAG-RFP retrovirus were infused into PV^Cre^; CAG^floxStopChR2-EYFP^ or SST^Cre^; CAG^floxStopChR2-EYFP^ mice to obtain GCs expressing RFP (RFP-GC), and PV- or SST-INs expressing ChR2 (ChR2-PV or ChR2-SST). Inversely, Ubi-ChR2-EGFP retrovirus were delivered into PV^Cre^; CAG^floxStoptd-Tomato^ or SST^Cre^; CAG^floxStoptd-Tomato^ to obtain GCs expressing ChR2 (ChR2-GC), and PV- or SST-INs expressing td-Tomato (Tom-PV or Tom-SST).

### Electrophysiological recordings

#### Slice preparation

Mice were anesthetized and decapitated at different weeks post injection (wpi) as indicated, and transverse slices were prepared as described previously (Marin-Burgin et al., 2012). Briefly, brains were removed into a chilled solution containing (in mM): 110 choline-Cl^−^, 2.5 KCl, 2.0 NaH2PO4, 25 NaHCO3, 0.5 CaCl2, 7 MgCl2, 20 glucose, 1.3 Na^+^-ascorbate, 0.6 Na^+^-pyruvate and 4 kynurenic acid. The hippocampus was dissected and transverse slices of septal pole (400 μm thick) were cut in a vibratome (Leica VT1200 S, Nussloch, Germany) and transferred to a chamber containing artificial cerebrospinal fluid (ACSF; in mM): 125 NaCl, 2.5 KCl, 2 NaH2PO4, 25 NaHCO3, 2 CaCl2, 1.3 MgCl2, 1.3 Na^+^-ascorbate, 3.1 Na^+^-pyruvate, and 10 glucose (315 mOsm). Slices were bubbled with 95% O2/5% CO2 and maintained at 30°C for at least 1 hour before experiments started.

#### Electrophysiology

Whole-cell and cell-attached recordings were performed at room temperature (23 ± 2 °C) using microelectrodes (4-6 MΩ for GCs and 3-5 MΩ for INs) filled with internal solution. All internal solution contained in common (in mM): 0.1 EGTA, 10 HEPES, 4 ATP-tris and 10 phosphocreatine, with pH 7.3 and 290 mOsm. To record INs or ChR2-GCs, we used internal solution with the following additional composition (in mM): 150 K-gluconate, 1 NaCl and 4 MgCl2. To measure IPSCs in RFP-GCs, we filled the recording electrodes with (in mM): 110 K-gluconate, 5 NaCl, 30 KCl and 4 MgCl2. Field recordings were performed using patch pipettes (2-4 MΩ) filled with 3 M NaCl. All recordings were obtained using Axopatch 200B amplifiers (Molecular Devices, Sunnyvale, CA), digitized (Digidata 1322A, Molecular Devices), and acquired at 10-20 KHz onto a personal computer using the pClamp 9 software (Molecular Devices).

Whole-cell voltage-clamp recordings were performed at a holding potential (Vh) of −70 mV, except for the experiment to study the reversal potential of GABAergic current onto GCs (Fig. 2). For GCs, series resistance was typically 10–20 MΩ, and experiments were discarded if higher than 25 MΩ. For INs, series resistance was typically 5–10 MΩ, and experiments were discarded if higher than 15 MΩ.

#### Recording target

Adult-born GCs expressing RFP or ChR2 were binned in the following age groups: 13-14 dpi (2 wpi), 20-22 dpi (3 wpi), 27-30 dpi (4 wpi), 40-44 dpi (6 wpi), 54-60 dpi (8 wpi) and 75-77 dpi (11 wpi). In previous work we have compared mature neurons born in 15-day-old embryos (which populate the outer granule cell layer), 7-day-old pups and adult mice, finding no functional differences among neuronal groups (Laplagne et al., 2006). Therefore, unlabeled neurons localized in the outer third of the granule cell layer were selected here as mature controls. Recorded neurons were visually identified in the granule cell layer by fluorescence (FITC fluorescence optics; DMLFS, Leica) and/or infrared DIC videomicroscopy. Criteria to include cells in the analysis were visual confirmation of fluorescent protein (RFP, Tom, GFP or YFP) in the pipette tip, attachment of the labeled soma to the pipette when suction is performed, and absolute leak current <100 pA and <250 pA at Vh for GCs and INs, respectively. Since INs are differentially distributed over distinct DG areas, we tried to maintain this proportion on the number of recorded INs in each region (Fig. 5; Fig. S1, Fig. S2).

#### Optogenetics

Patch-clamp recordings were carried out in GCs or in DG INs from hippocampal slices containing several INs or GCs expressing ChR2 (ChR2-PVs, ChR2-SSTs or ChR2-GCs). The latter were visualized by their EGFP or EYFP expression, as previously described (Toni et al., 2008). ChR2-neurons were stimulated using a 447 nm laser source delivered through the epifluorescence pathway of the upright microscope (FITC filter, 63X objective for whole-cell recordings, and 20X for field recordings) commanded by the acquisition software. Laser pulses (1 ms onto ChR2-GCs and 0.2 ms onto ChR2-INs) were delivered at 0.07 Hz while postsynaptic currents were recorded in voltage-clamp configuration. The laser power intensity was <150 mW. EPSCs onto INs were isolated by voltage clamping the neurons at the reversal potential of the IPSC (Vh = −70 mV). When analyze the spikes evoked onto INs through GC-ChR2 stimulation or activation of afferent pathway, the former were hold at −60 mV. To study unitary IPSCs, laser intensity was lowered to reach a condition where the GCs displayed both failures (in at least 10% of the total trials) and small IPSCs. Glutamatergic currents were blocked by KYN 4 mM and GABAergic currents were blocked by PTX 100 μm.

#### Field recordings

Medial perforant path (mPP) stimulation was performed by placing a steel monopolar electrode in the middle of the molecular layer, and current pulses ranging from 10 to 150 μA (0.2 ms) were applied at 0.07 Hz. The recording microelectrode was placed in the GCL to record the population spike (pop-spike) in response to mPP stimulation (Marin-Burgin et al., 2012). Experiments were performed at stimulus intensities that evoked 30 - 55 % of maximal pop-spike amplitude. Population activity was recorded by several subsequent trials until stable pop-spike amplitude was obtained. At that moment, a laser pulse (0.2 ms) was paired to mPP stimulation at different times (as indicated), alternating 5 consecutives trials with the laser on and 5 trials off.

#### Reversal potential of GABAergic currents onto GCs

Outer molecular layer (OML) stimulation was performed by placing a steel monopolar electrode in the outer third of the molecular layer, at least 300 μm away from the recording site. Current pulses ranging from 40 to 100 μA (0.2 ms) were applied at 0.05 Hz to recruit GABAergic current of dendritic origin. In addition, IPSCs evoked onto GCs in response to optogenetics stimulation of ChR2-INs were measured. This study was performed in presence of kynurenic acid.

### In vivo assays. EE Exposure

Two weeks after RV infusion, mice were exposed for two weeks to an EE consisting of a large cage (80 cm × 40 cm × 20 cm) containing tunnels of different lengths, toys, and two running wheels. The location of the objects in the EE were changed after a week of exposure. Control mice were left in a regular cage with two running wheels (consistent with our experiments). At 4 wpi, animals were prepared for electrophysiological recordings

### Immunofluorescence

Immunostaining was performed in 60 μm free-floating coronal sections throughout the brain from six weeks old PV^Cre^ and SST^Cre^; CAG^FloxStopChR2^ mice. Antibodies were applied in TBS with 3% donkey serum and 0.25% Triton X-100. Triple labeled immunofluorescence was performed using the following primary antibodies: GFP (Green Fluorescent Protein, Chicken antibody IgY Fraction 1:500, Aves Labs Inc.), PV (mouse anti-Parvalbumin monoclonal antibody, 1:3000, Swant) and SOM (rat-anti Somatostatin monoclonal antibody 1:250, Millipore). The following corresponding secondary antibodies were used: donkey anti-chicken Cy2, donkey anti-mouse Cy5 and donkey anti-rat Cy3, (1:250; Jackson ImmunoResearch Laboratories). Incubation with Dapi (10 minutes) was applied to avoid fluorescence bleaching when slice characterization was performed.

### Confocal microscopy

Sections from the hippocampus (antero-posterior, −0.94 to −3.4 mm from bregma) according to the mouse brain atlas (Paxinos and Franklin, 2004) were included. Images were acquired using Zeiss LSM 510 Meta microscope (Carl Zeiss, Jena, Germany). Analysis of antibody expression was restricted to cells with fluorescence intensity levels that enabled clear identification of their somata. Images were acquired (40X, NA 1.3, oil-immersion) and colocalization for the three markers was assessed in z-stacks using multiple planes for each cell. Colocalization was defined as positive if all markers were found in the same focal plane.

### Data analysis

Analysis of all recordings was performed off-line using in-house made Matlab routines.

#### Intrinsic Properties

Membrane capacitance and input resistance were obtained from current traces evoked by a hyperpolarizing step (10 mV, 100 ms). Spiking profile was recorded in current-clamp configuration (membrane potential was kept at −70 mV by passing a holding current) and the threshold current for spiking was assessed by successive depolarizing current steps (10 pA for GCs and 50 pA for INs; 500 ms) to drive the membrane potential (Vm) from resting to 0 mV.

Action potential threshold was defined as the point at which the derivative of the membrane potential dVm/dt was 5 mV/ms (data not shown). AP amplitude was measured from threshold to positive peak and after-hyperpolarization amplitude, from threshold to negative peak during repolarization. Time between consecutive spikes (interspike interval, ISI) was measured from peak to peak. Instantaneous frequency was calculated from ISI and adaptation ratio was defined as the ISI ratio between the third spike and the last spike. To perform the whole spiking characterization, we measured the threshold current intensity and a stimulus intensity three times higher than the threshold was used to evaluate all the parameters.

#### Postsynaptic Currents

Statistical methods were used to differentiate laser-responsive cells and laser-evoked events from spontaneous activity using in-house Matlab routines. Events were identified as peaks in the low-pass filtered current (<250 Hz) when exceeded 4 standard deviations of the noise level (measured at >500 Hz high-pass filtered current). The onset of an event was defined as the time in which 10 % of the maximum amplitude was reached in the unfiltered signal. Once all events were identified, a cell was classified as responsive to laser stimulation if there was a tendency greater than chance for events to accumulate within a time window of 12 ms after laser stimulation (p < 0.05). In order to achieve such a classification, the probability distribution of a similar accumulation of spontaneous events happening by pure chance was determined for each cell using a 2000 step shuffling procedure. Once a cell was classified as responsive to laser, spontaneous and laser-evoked events were differentiated.

In all cases, reported PSCs values for peak amplitude correspond to the product of the mean value for positive trials and the probability of success, taken as the fraction of trials in which an evoked response was observed. The rise time was calculated from 20% to 80% (EPSC) or 70% (IPSC) of peak amplitude, and decay time was calculated from 80% (EPSC) or 70% (IPSC) to 30%.

#### Response to repetitive stimulation

The charge of laser evoked events during repetitive stimulation was measured within a time window equal to the distance between two consecutive laser pulses, starting at the corresponding pulse. To analyze short-term plasticity, we calculated the charge ratio during repetitive stimulation. To perform this normalization, we used the response evoked by the first pulse for INs onto GCs synapses and the charge related to the last pulse for GCs onto INs synapses.

### Statistical analysis

Unless otherwise specified, data are presented as mean ± SEM. Normality was assessed using Shapiro-Wilk’s test, D’Agostino & Pearson omnibus test, and Kolmogórov-Smirnov’s test, at a significance level of 0.05. A distribution was considered as normal if all tests were passed. When a data set did not satisfy normality criteria, non-parametric statistics were applied. Two-tailed Mann-Whitney’s test was used for single comparisons, two-tailed Wilcoxon matched pairs signed rank test was applied for paired values, Kruskal-Wallis test by ranks was employed to compare multiple unmatched groups and Friedman test followed by Dunn’s post test was used to compare multiple matched groups. For normal distributions, homoscedasticity was assessed using Bartlett’s test and F-test, at a significance level of 0.05. For homogeneous variances, two-tailed t-test was used for single comparisons, and ANOVA test followed by post-hoc Bonferroni’s test was used for multiple comparisons. Two sample Kolmogorov-Smirnov test was applied to compare cumulative distributions. Two-tailed Fisher’s exact test (small sample size) or Chi-square test were used in the analysis of contingency tables.

## SUPPLEMENTAL FIGURES

**Figure S1. Characterization of dentate gyrus PV-INs.** (A) (left) Experimental scheme showing whole-cell recordings on Tom-PVs. (Right) PV-IN recording displaying high frequency firing in response to 600 pA-step. Scale bar: 100ms, 20mV. Spiking profile was assessed using the following measures: action potential (AP) amplitude (B), after hyperpolarization amplitude (C), AP width (D), firing maximum frequency (E), and AP adaptation ratio (F). Membrane passive properties are described by resting membrane potential (G), input resistance (H) and membrane capacitance (I). Sample sizes were 11-23 cells in 5-9 animals. Statistical comparisons were done using Kruskal-Wallis test. (J) Recorded Tom-PVs were characterized by their location in the dentate gyrus, divided into three main areas: hilus, molecular layer (ML) and granular cell layer (GCL). The spatial distribution of recorded Tom-PVs is not significantly different among groups. The total numbers of cells are shown on top of each column and the numbers of interneurons in each area are displayed within the columns. Statistical comparisons were done using Chi-square test (p=0.819).

**Figure S2. Characterization of dentate gyrus SST-INs.** (A) Experimental scheme for whole-cell recordings on Tom-SSTs. (B) SST-IN recording showing repetitive firing in response to 510 pA step. We observed four different spiking profiles: fast spiking (FS), accomodating (Ac), stuttering (St) and single-spike (SS). Proportions (%) corresponding to each spiking profile are indicated. Scale bar: 100ms, 20mV. (C) Percentage of SST-INs spiking profiles recorded at different GC stages. Membrane passive properties are described by input resistance (D) and membrane resting potential (E). Passive properties were then classified for each spiking profile: input resistance (F) and membrane resting potential (G). (H) Recorded SST-INs were characterized by their location in the dentate gyrus. We used three main areas: hilus, molecular layer (ML) and granular cell layer (GCL). The spatial distribution of recorded SST-IN is not significantly different among groups. Total numbers of cells are shown on top of each column and numbers of interneurons in each area are displayed within the columns test. Statistical comparisons were done using Chi-square test (p=0.748). (I) The spatial distribution for INs presenting each spiking profile is similar except for FS, which is mainly located in the GCL.

**Figure S3. In-depth characterization of IPSCs evoked by ChR2-PV activation.** (A) PV-INs spiking elicited by optogenetics. (i) Experimental scheme shows PV-INs recording after brief laser pulses. (ii) Cell attached recording showing PV-INs reliable spiking after single laser pulse stimulation (0.2 ms, 0.07 Hz, blue marks). Scale bar, 20 pA, 5 ms. (B) Total number of spikes per single pulse when applying saturating intensity of laser stimulation. All measured ChR2-PVs were responsive to a single-pulse laser stimulation. N = 13 cells / 9 mice. (C) Time to onset of the first PV spike response. (D) Experimental scheme for RFP-GC recording after PV-INs stimulation. (E) IPSCs elicited by laser pulses (0.2 ms, blue marks) delivered at low frequency (0.07 Hz), recorded from GCs at the indicated ages. Traces depict individual sweeps (gray) and their average (black). Scale bars, 100 pA, 10 ms. (F) Normalized traces highlighting the differences in kinetics for responses recorded from GCs at 3 and 8 wpi. Scale bar, 10 ms, 0.2 au. (G) Percentage of adult-born GCs presenting response to activation of PV-INs. Total amount of GCs recorded are shown on top of each column, and the numbers of responsive GCs are displayed within columns. Statistical comparisons were done using Fisher’s exact test. We measured the time to onset (H), half-width (I) and the decay time (J) for IPSC elicited by PV-INs. Statistical comparisons were done using one-way ANOVA followed by *post hoc* Bonferroni’s test for multiple comparisons against mature condition. (K, L) Representative histogram of IPSC amplitude evoked by minimal stimulation for GCs at 4 wpi (K) and mature (L). Insets show normalized average traces in response to minimal (IPSC-unit) and maximal (IPSC-sat) stimulation. Scale bars, 10 ms, 0.1 au. (M) The number of IPSC-units contained in IPSC-sat are not different among immature and mature GCs. Black circles correspond to the examples shown in *L-M*. N = 10 (4 wpi) and 10 (mature) cells. Statistical comparison was done using two-tailed t-test (p=0.128). *p*<0.05 (*), *p*<0.01 (**), *p*<0.001 (***).

**Figure S4. In-depth characterization of IPSCs evoked by ChR2-SST activation.** (A) SST-INs spiking elicited by optogenetic. (i) Experimental scheme shows SST-INs recording after laser stimulation. (ii) Representative cell attached recordings showing SST spiking after single laser-pulse stimulation (0.2 ms, 0.07 Hz, blue marks). Scale bar, 20 pA, 5 ms. (B) Total number of spikes per single pulse. All measured ChR2-SSTs were responsive to a single-pulse laser stimulation. N = 8 cells / 5 mice. (C) Time to onset of the first SST-IN spike. (D) Simplified experimental schematic shows RFP-GCs recording when ChR2-SSTs are stimulated. (E) Coefficient of variation for proximal and distal IPSC amplitudes recorded in adult-born GCs. Statistical comparisons were done using two-way ANOVA followed by *post hoc* Bonferroni’s test for multiple comparisons. (F, G) Percentage of adult-born GCs presenting response to activation of SST-INs, for both proximal (F) and distal (G) IPSCs. Total number of recorded GCs are shown on top of each column, and numbers of responsive and non-responsive GCs are displayed within columns. Statistical comparisons were done using Fisher’s exact test. (H-K) Proximal IPSC kinetic. (H) Normalized traces highlighting the differences in kinetics for responses evoked in adult-born GCs at 3 and 8 wpi. Scale bars, 10 ms, 0.2 norm. We measured time to onset (I), half-width (J) and decay time (K). Statistical comparisons were done using one-way ANOVA followed by *post hoc* Bonferroni’s test for multiple comparisons against mature condition. (L-O) Distal IPSC kinetic, corresponding to *H-K*. *p*<0.01 (**), *p*<0.001 (***).

**Figure S5. In-depth characterization of EPSCs evoked onto PV- and SST-INs by ChR2-GC activation.** (A) ChR2-GCs spiking elicited by optogenetics. (i) Experimental scheme of GCs recording after optogenetic stimulation. (ii) Representative cell-attached recordings in ChR2-GCs show reliable spiking evoked by brief laser pulse (1ms, 0.07Hz, blue marks). Representative data from 4 wpi GCs. Scale bar, 10 ms, 50 pA. Sample sizes were 10-19 neurons in 5-9 mice. (B) Total number of spikes per single pulse at each GCs stages. Gray dots correspond to single neurons. (C) Time to onset of the first spike elicited on ChR2-GCs by single laser-pulse stimulation. Activation of adult-born GCs elicits EPSCs onto PV-INs (D-J) and SST-INs (K-Q). (D) Experimental scheme depicting laser-activated GCs and PV-IN recording. (E) Recordings of PV-INs EPSCs elicited by laser-pulse stimulation of adult-born GCs at 4 and 8 wpi. Traces depict all sweeps in the experiment (gray) and their average (black). Scale bar, 10 ms, 50 pA. EPSC peak amplitude (F), rise time (G), time to onset (H), half-width (I) and decay time (J) are presented. Sample sizes were 11-23 neurons in 5-9 mice. (K) Experimental scheme for SST-INs recording. (L) Recordings of SST-INs EPSCs elicited by GCs activation at 6 and 8 wpi. Scale bar, 10 ms, 50 pA (top), 25 pA (bottom). (M-Q) Corresponding to *F-J* for SST-INs recordings. Sample sizes were 12-17 neurons in 2-7 mice. Hollow symbols correspond to example traces. Statistical comparisons were done using Kruskal-Wallis test followed by *post hoc* Dunn’s multiple comparisons. *p*<0.05 (*) and *p*<0.01 (**).

