## Supplemental data for "Differential coupling of adult-born granule cells to parvalbumin and somatostatin interneurons"

**Figure S5. In-depth characterization of EPSCs evoked onto PV- and SST-INs by ChR2-GC activation.** (A) ChR2-GCs spiking elicited by optogenetics. (i) Experimental scheme of GCs recording after optogenetic stimulation. (ii) Representative cell-attached recordings in ChR2-GCs show reliable spiking evoked by brief laser pulse (1ms, 0.07Hz, blue marks). Representative data from 4 wpi GCs. Scale bar, 10 ms, 50 pA. Sample sizes were 10-19 neurons in 5-9 mice. (B) Total number of spikes per single pulse at each GCs stages. Gray dots correspond to single neurons. (C) Time to onset of the first spike elicited on ChR2-GCs by single laser-pulse stimulation. Activation of adult-born GCs elicits EPSCs onto PV-INs (D-J) and SST-INs (K-Q). (D) Experimental scheme depicting laser-activated GCs and PV-IN recording. (E) Recordings of PV-INs EPSCs elicited by laser-pulse stimulation of adult-born GCs at 4 and 8 wpi. Traces depict all sweeps in the experiment (gray) and their average (black). Scale bar, 10 ms, 50 pA. EPSC peak amplitude (F), rise time (G), time to onset (H), half-width (I) and decay time (J) are presented. Sample sizes were 11-23 neurons in 5-9 mice. (K) Experimental scheme for SST-INs recording. (L) Recordings of SST-INs EPSCs elicited by GCs activation at 6 and 8 wpi. Scale bar, 10 ms, 50 pA (top), 25 pA (bottom). (M-Q) Corresponding to *F-J* for SST-INs recordings. Sample sizes were 12-17 neurons in 2-7 mice. Hollow symbols correspond to example traces. Statistical comparisons were done using Kruskal-Wallis test followed by *post hoc* Dunn's multiple comparisons. $p<0.05$ (*) and $p<0.01$ (**).

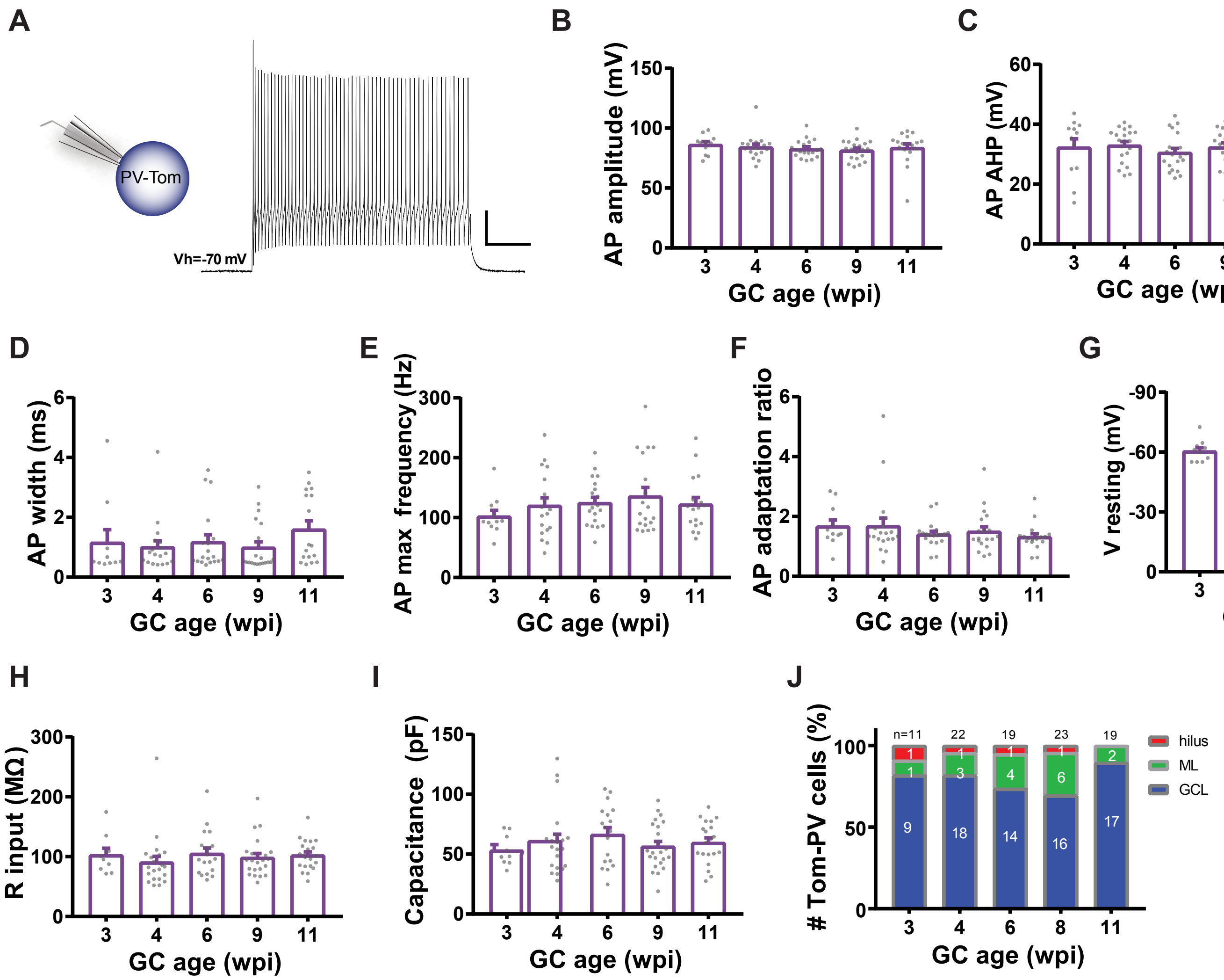
A
B
C
D
E
F
G
H
I
J
PV-Tom
Vh=-70 mV
AP amplitude (mV)
AP AHP (mV)
AP width (ms)
AP max frequency (Hz)
AP adaptation ratio
V resting (mV)
R input (MΩ)
Capacitance (pF)
### Tom-PV cells (%)
GC age (wpi)
n=11
22
19
23
19
hilus
ML
GCL

Fig. S1

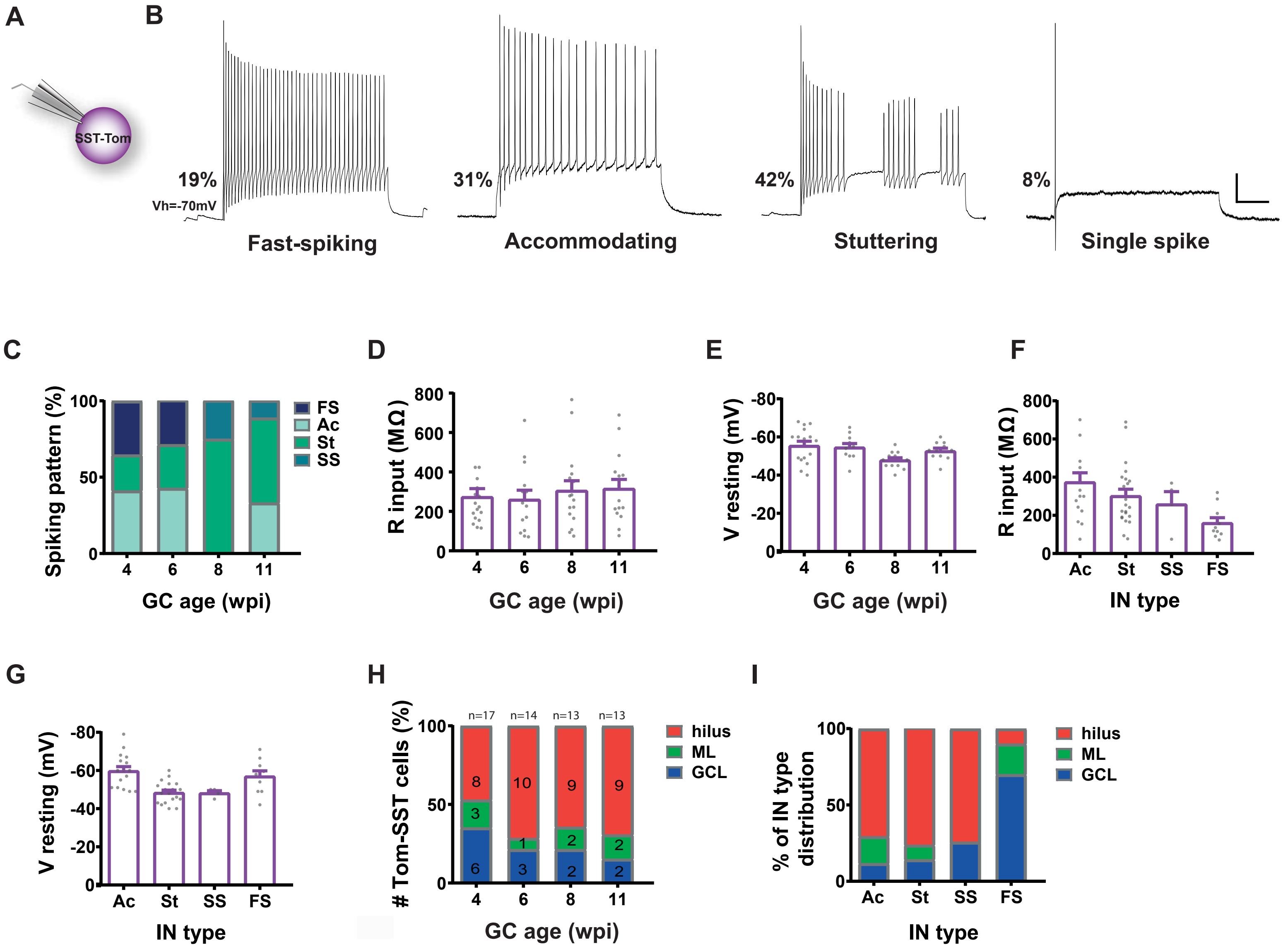
A
B
SST-Tom
19%
Vh=-70mV
Fast-spiking
31%
Accommodating
42%
Stuttering
8%
Single spike
C
Spiking pattern (%)
100
50
0
4
6
8
11
GC age (wpi)
FS
Ac
St
SS
D
R input (MΩ)
800
600
400
200
0
4
6
8
11
GC age (wpi)
E
V resting (mV)
-80
-60
-40
-20
0
4
6
8
11
GC age (wpi)
F
R input (MΩ)
800
600
400
200
0
Ac
St
SS
FS
IN type
G
V resting (mV)
-80
-60
-40
-20
0
Ac
St
SS
FS
IN type
H
### Tom-SST cells (%)
n=17
n=14
n=13
n=13
100
50
0
8
3
6
10
1
3
9
2
2
9
2
2
4
6
8
11
GC age (wpi)
hilus
ML
GCL
I
% of IN type distribution
100
50
0
Ac
St
SS
FS
IN type
hilus
ML
GCL

Fig. S2

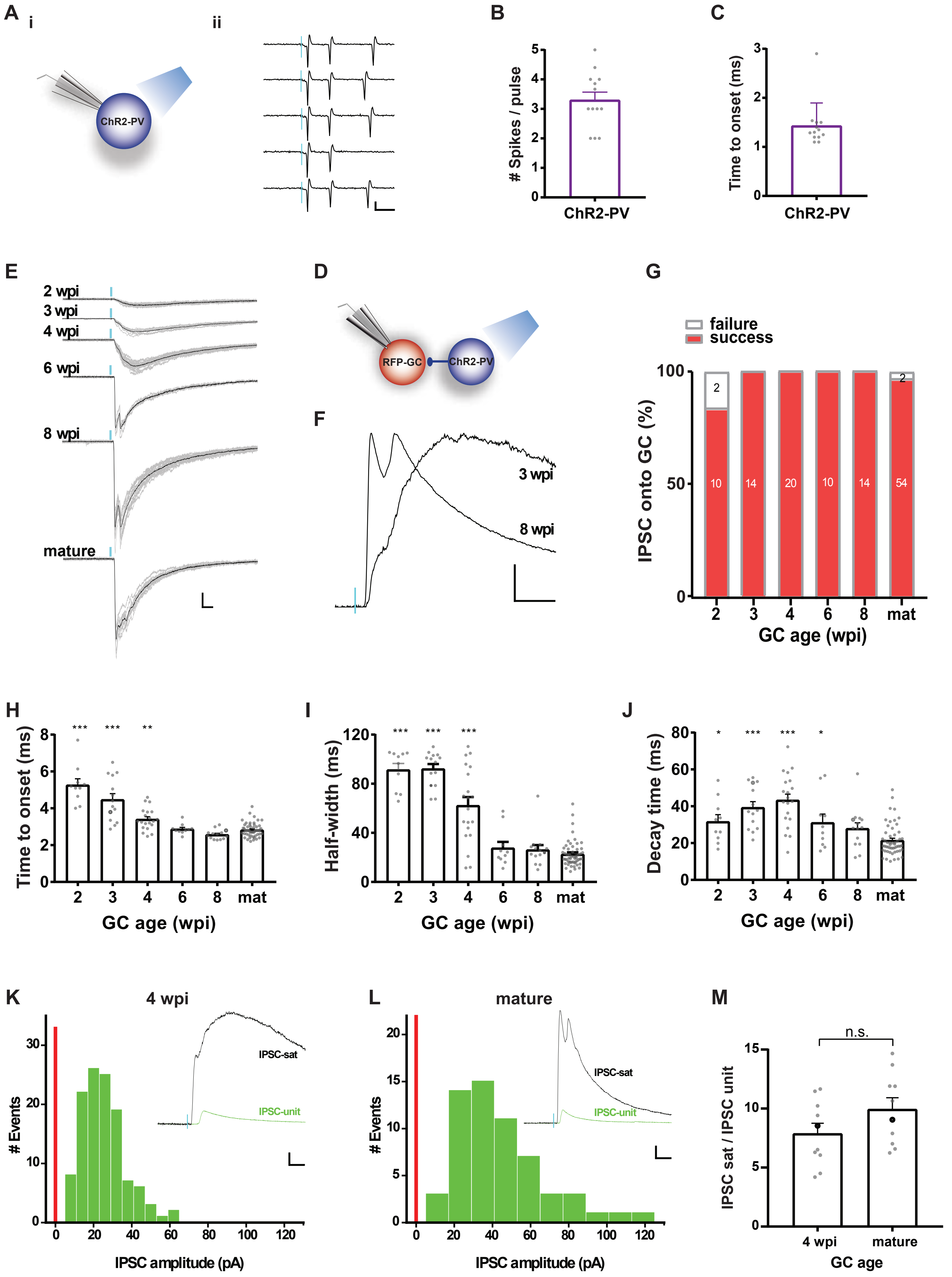
A
i
ii
ChR2-PV
B
### Spikes / pulse
ChR2-PV
C
Time to onset (ms)
ChR2-PV
E
2 wpi
3 wpi
4 wpi
6 wpi
8 wpi
mature
D
RFP-GC
ChR2-PV
F
3 wpi
8 wpi
G
failure
success
IPSC onto GC (%)
GC age (wpi)
H
Time to onset (ms)
GC age (wpi)
I
Half-width (ms)
GC age (wpi)
J
Decay time (ms)
GC age (wpi)
K
4 wpi
IPSC-sat
IPSC-unit
### Events
IPSC amplitude (pA)
L
mature
IPSC-sat
IPSC-unit
### Events
IPSC amplitude (pA)
M
n.s.
IPSC sat / IPSC unit
4 wpi
mature
GC age

Fig. S3

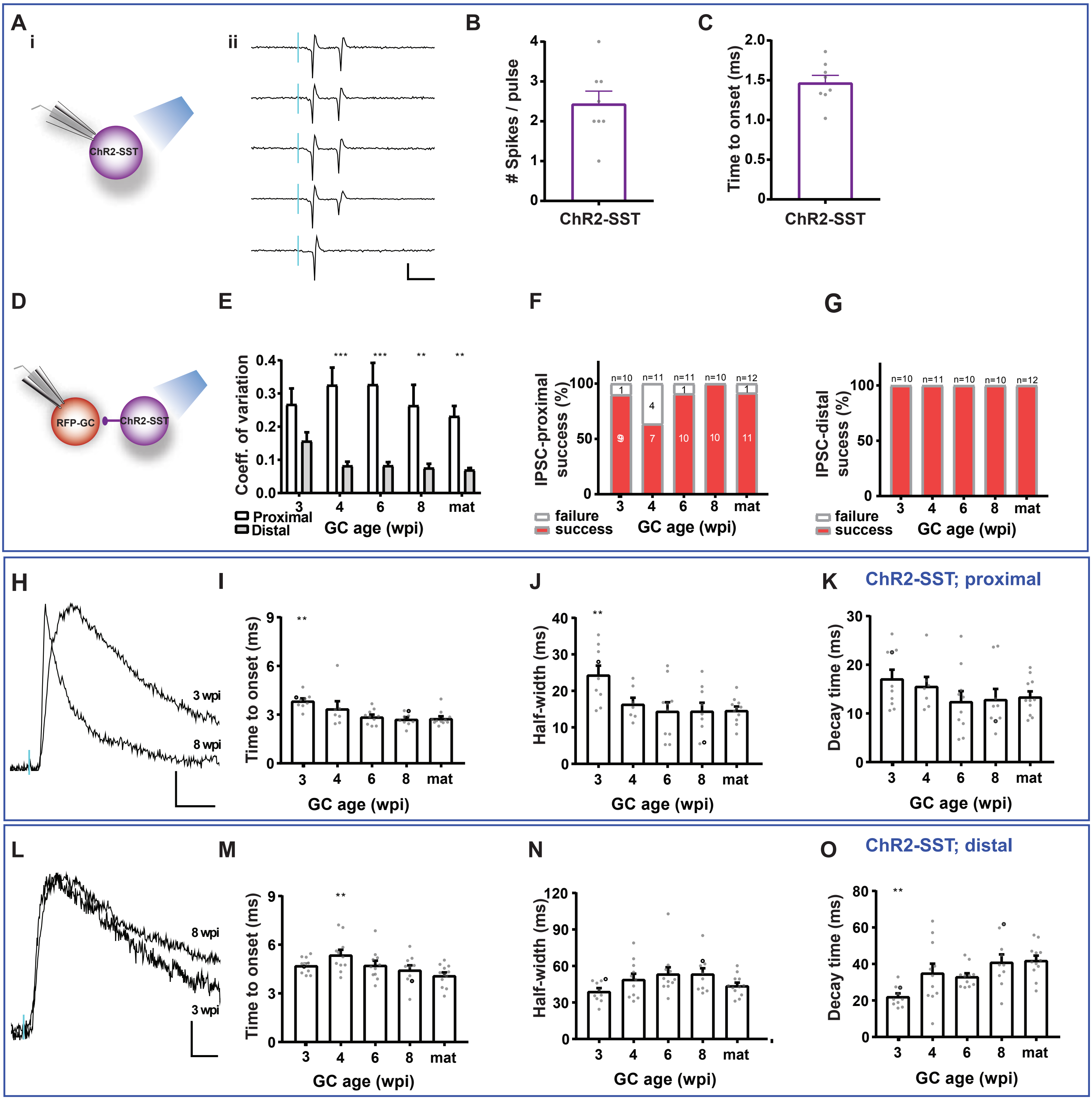
A
i
ii
ChR2-SST
B
### Spikes / pulse
ChR2-SST
C
Time to onset (ms)
ChR2-SST
D
RFP-GC
ChR2-SST
E
Coeff. of variation
Proximal
Distal
GC age (wpi)
F
IPSC-proximal success (%)
failure
success
GC age (wpi)
G
IPSC-distal sucess (%)
failure
success
GC age (wpi)
H
3 wpi
8 wpi
I
Time to onset (ms)
GC age (wpi)
J
Half-width (ms)
GC age (wpi)
K
ChR2-SST; proximal
Decay time (ms)
GC age (wpi)
L
8 wpi
3 wpi
M
Time to onset (ms)
GC age (wpi)
N
Half-width (ms)
GC age (wpi)
O
ChR2-SST; distal
Decay time (ms)
GC age (wpi)

Fig. S4

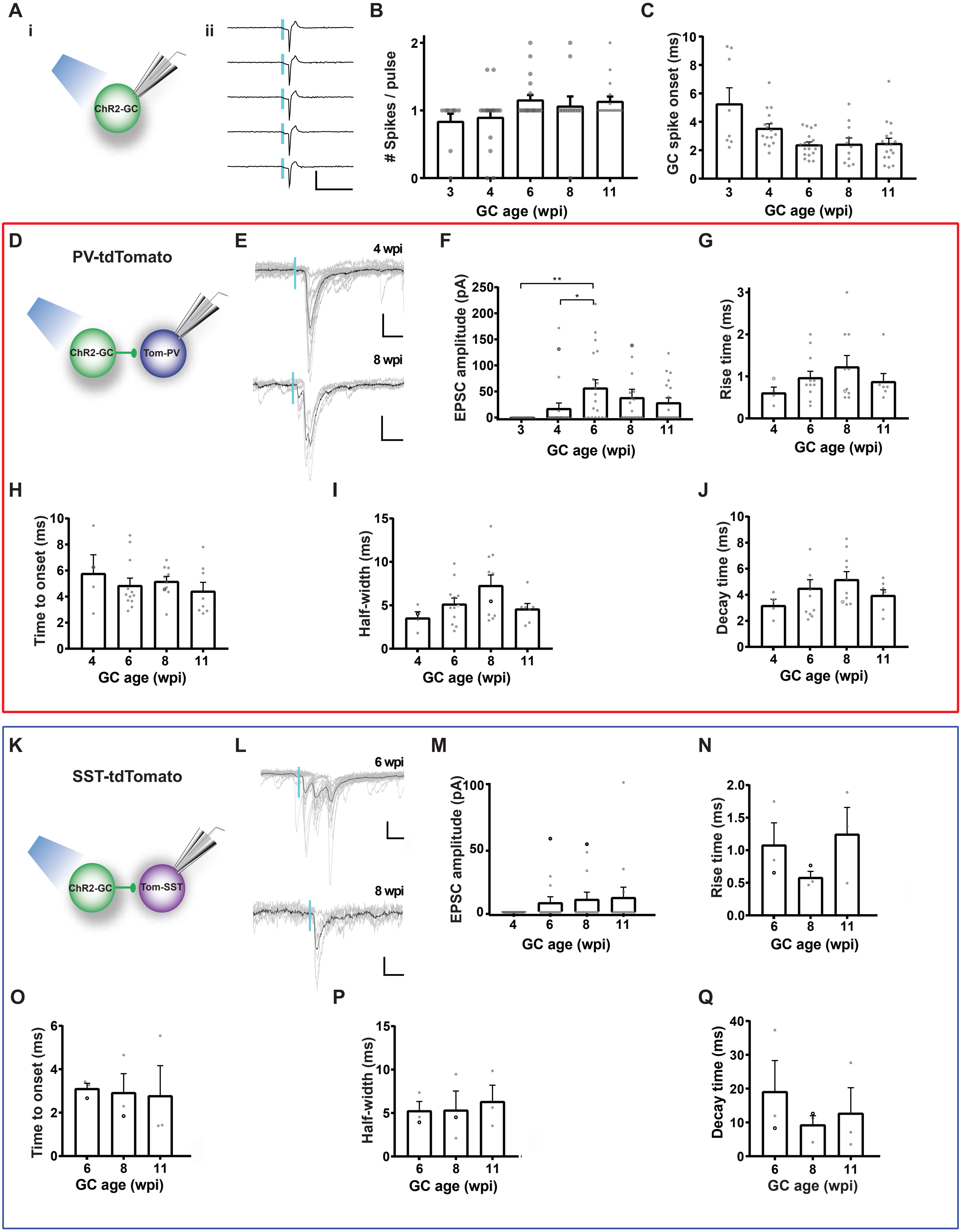
A
i
ii
ChR2-GC
B
### Spikes / pulse
GC age (wpi)
C
GC spike onset (ms)
GC age (wpi)
D
PV-tdTomato
ChR2-GC
Tom-PV
E
4 wpi
8 wpi
F
EPSC amplitude (pA)
GC age (wpi)
G
Rise time (ms)
GC age (wpi)
H
Time to onset (ms)
GC age (wpi)
I
Half-width (ms)
GC age (wpi)
J
Decay time (ms)
GC age (wpi)
K
SST-tdTomato
ChR2-GC
Tom-SST
L
6 wpi
8 wpi
M
EPSC amplitude (pA)
GC age (wpi)
N
Rise time (ms)
GC age (wpi)
O
Time to onset (ms)
GC age (wpi)
P
Half-width (ms)
GC age (wpi)
Q
Decay time (ms)
GC age (wpi)

Fig. S5
